# Pitfalls of machine learning models for protein-protein interactions

**DOI:** 10.1101/2022.02.07.479382

**Authors:** Loïc Lannelongue, Michael Inouye

## Abstract

Protein-protein interactions (PPIs) are essential to understanding biological pathways as well as their roles in development and disease. Computational tools, based on classic machine learning, have been successful at predicting PPIs *in silico*, but the lack of consistent and reliable frameworks for this task has led to network models that are difficult to compare and discrepancies between algorithms that remain unexplained. To better understand the underlying inference mechanisms that underpin these models, we designed an open-source framework for benchmarking that accounts for a range of biological and statistical pitfalls while facilitating reproducibility. We use it to shed light on the impact of network topology and how different algorithms deal with highly connected proteins. By studying functional genomics-based and sequence-based models on human PPIs, we show their complementarity as the former performs best on lone proteins while the latter specialises in interactions involving hubs. We also show that algorithm design has little impact on performance with functional genomic data. We replicate our results between both human and *S. cerevisiae* data and demonstrate that models using functional genomics are better suited to PPI prediction across species. With rapidly increasing amounts of sequence and functional genomics data, our study provides a principled foundation for future construction, comparison and application of PPI networks.

## Introduction

Protein-protein interactions (PPIs) are central to protein function and inform a wide range of biomedical applications, from mechanistic studies [1], [2] to drug development [3], [4]. Better understanding these interactions is critical for successfully mapping biological pathways, but the diversity of PPIs and the scale of the network make this a difficult task. Experimental methods to map PPIs exist, but, despite progress in systematic mapping [5], [6], even high-throughput ones are not yet able to determine all possible interactions.

Through generalisable patterns learned on varied interaction data, computational methods can complement experiments by addressing the issue of scalability and measurement bias. Given a pair of proteins and some characteristics of each one, machine learning models can learn to predict the likelihood of interaction. Numerous methods have been developed for this, using the full range of machine learning models, from early work on *Saccharomyces cerevisiae* [7]–[10] to algorithms dedicated to human PPIs [11]–[15]. Yet, despite a wealth of tools, the mechanics and consequences of the underlying inference are still poorly understood, and it is unclear why models with similar performance make vastly different predictions. Reported performance scores often cannot be compared or replicated due to proprietary data and inconsistent or flawed assessment methods [16], [17]. This prompted recent efforts to benchmark published PPI prediction models more rigorously using common datasets and testing strategies [16], [18]. These benchmarking studies hinted at some fundamental differences in how algorithms predict PPIs, e.g. some sequence-based models mostly leverage biases in the data [16], which highlighted the need for inference mechanisms to be investigated further and with the same rigour.

In the absence of such studies, *in silico* PPI analyses are difficult to reconcile, the development of new models is inefficient, follow-up mechanisms studies are likely undermined and, ultimately, there are different versions of the underlying molecular networks that describe protein function.

A unified framework to understand PPI algorithms would improve the development and reliable assessment of new models, and would facilitate the overdue widespread adoption of PPI predictions for downstream analysis. Replicable, trustworthy and generalisable high-performing models can capture more causal biology and enhance many aspects of biological research such as experimental designs and drug development.

In this work, we study and compare two of the main building blocks of PPI prediction algorithms, based on functional genomic (FG) information or amino acid sequences alone. We highlight why both perspectives are still relevant today and how each adapts to the PPI network’s topology. In particular, we show that the presence of highly connected proteins in the networks has a drastic impact on prediction models and is an area where FG and sequence models diverge. We also replicate these results between human and yeast (*S. cerevisiae*) and show how each tool performs on cross-species predictions. To elucidate the prediction mechanisms of these models, we design a robust and standardised approach to investigate *in silico* PPI prediction tools that accounts for both biological and statistical pitfalls and leverages the strength of large, open-source and professionally curated databases. We make publicly available benchmarking datasets for human and yeast PPIs to accelerate future discoveries and lay the foundations for similar datasets for other organisms. This work gives critical insight into each approach’s strengths and weaknesses and provides robust foundations for future developments in PPI prediction models.

## Results

### A robust and open-source benchmarking pipeline to understand PPI prediction models

The lack of a consistent way to investigate PPI prediction algorithms has hindered their development and reduced their impact by making it difficult to reuse models for downstream analyses [16], [19]. Benchmarks are important for replicability [20], and when combined with carefully curated datasets, they enable fast development through trial and error. Our Benchmarking Pipeline for the Prediction of Protein-Protein Interactions in Humans (B4PPI-Human) includes both carefully selected training and testing sets and a collection of input features to enable such trials. Standard UniProt IDs are used throughout to easily combine these with other data sources. Relevant metrics are selected with guidelines on how to share them. All this, alongside the pre-processing steps and relevant guidelines, is made available online and can be downloaded easily from https://github.com/Llannelongue/B4PPI. An example of a reporting sheet is in Figure 1.

**Figure 1:**
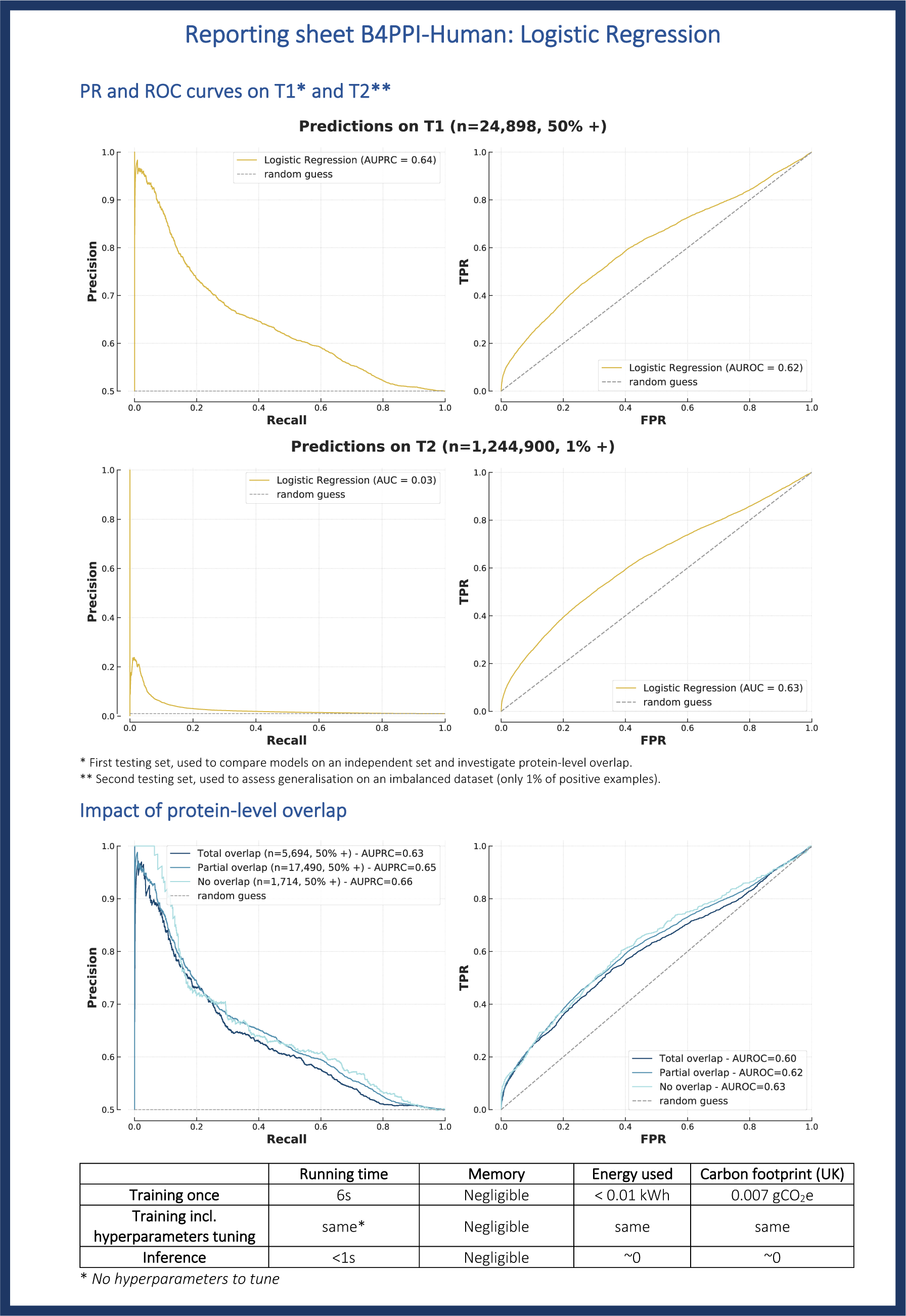
Reported performance sheet of the logistic regression (FG-based) on B4PPI-Human.

The complexity of the underlying biological mechanisms of PPIs introduces pitfalls that need to be considered when evaluating models. First, the way non-interacting proteins are selected for training is important. While some efforts have used proteins known to be localised in different parts of the cell [7], [14], [21], [22], this has been shown to be unreliable and a source of significant bias that overestimates accuracy [23]. An alternative is to use a database of experimentally tested non-interacting proteins, but leading resources such as Negatome have only ∼1,300 pairs and thus offer limited coverage [21], [24]. Considering the scarcity of PPIs, randomly sampling pairs of proteins has a very low risk of false negative and limits sampling bias (i.e. focusing on well-studied proteins) [23], [25]. However, the impact of the associated imbalance between interacting and non-interacting proteins should be taken into account when training models on balanced datasets [26]. Lastly, each observation is itself a pair of proteins. Even when ensuring that the two sets don’t have pairs in common, there can be individual proteins present in both the training and testing sets. This protein-level overlap, often overseen, has been shown to significantly affect the performance of an algorithm and should therefore be properly assessed [27]. Despite being documented in the literature, these pitfalls are still unevenly accounted for in published works [16]. This, alongside inconsistencies in the choice of testing sets and performance metrics explains why, despite the number of algorithms released in recent years, the underlying inference mechanisms are still not well understood.

The essential aspects of training and assessment that should be systematically accounted for are (1) the quality of the positive examples (i.e. the interacting proteins), (2) how non-interacting proteins are selected for the gold standard, (3) a suitable split between training and testing sets, in particular regarding individual proteins, and (4) the metrics to evaluate and compare models. B4PPI seeks to address these four aspects of benchmarking.

When building a gold standard for machine learning algorithms, quality and representativity are the most important aspects to consider, which makes IntAct [28] a database of choice for interacting proteins. It aggregates reliable evidence of molecular interactions from over 20,000 publications, which are manually curated, and includes data from other interactions databases such as the members of the IMEx consortium [29]. Besides, by aggregating PPIs obtained from different experiments, each with different technical limitations, IntAct limits measurement bias. We further limited the risk of false positives by removing low-quality interactions, for example the ones based on spatial colocalisation only (Methods). The final dataset comprised 78,229 interactions, covering 12,026 proteins (out of the 20,386 registered in UniProt).

To select non-interacting proteins to serve as negative examples, randomly sampling protein pairs is the approach with the lowest probability of error considering the scarcity of the PPI network [30]. Non-interacting proteins can be sampled using a uniform distribution, i.e. all proteins have an equal probability of being selected, which leads to an unbiased set, representative of the general population of protein pairs. However, PPI networks are known to be similar to scale-free networks, i.e. composed of a few highly connected nodes, called *hubs*, and numerous *lone* proteins with few interactions [31] (Supplementary Figure 1). Consequently, hubs are over-represented in a set of PPIs. For example in our curated set from IntAct, the top 20% of proteins by number of interactions were involved in 94% of PPIs. But when uniformly sampling protein pairs, the same top 20% were only involved in 37% of non-interacting proteins. Although expected, this can be an issue for machine learning algorithms that would identify hubs and systematically predict a positive interaction when hubs are involved. Such a strategy would maximise accuracy on the training set but lead to a majority of false positives when making predictions on new pairs. To mitigate this, a balanced sampling can be used [32], where the probability of sampling a protein for the negative set is proportionate to its frequency in the positive set. It has been shown that each strategy serves a different purpose [25]; balanced sampling is beneficial for training models but shouldn’t be used for evaluating them, as the induced bias makes metrics less meaningful. This was the strategy implemented here, where non-interacting proteins were selected with balanced sampling for the training set and uniform sampling for the testing sets.

In the presence of limited data, the division of the gold standard between training and testing sets is critical to simultaneously optimise learning and obtain meaningful generalisation metrics. Here, the testing set should achieve several objectives, (1) provide performance metrics on a new, independent set, (2) measure the impact, or absence of impact, of protein-level overlap, (3) demonstrate how the model can generalise to real-world data. Since a single set cannot achieve simultaneously (2) and (3), as the careful selection of proteins to measure overlap biases the dataset, we designed two testing sets *T1* and *T2* (Methods). *T1* should be used to compare different approaches with an independent set and investigate protein-level overlap, and *T2* should be used to assess generalisation. *T1* was built by purposefully leaving some proteins out of the training set; we demonstrated the importance of this as dividing the training and testing sets conventionally (using, for example, the popular *scikit-learn* library) resulted in almost all pairs (95%) in the testing set sharing at least one protein with the training set (Figure 2), which may lead to overestimating performances [27]. *T2*, with only 1% of positive examples (Supplementary Table 1), can then be used to assess how models perform in a more realistic setting where positive interactions are rare compared to non-interacting proteins.

**Figure 2:**
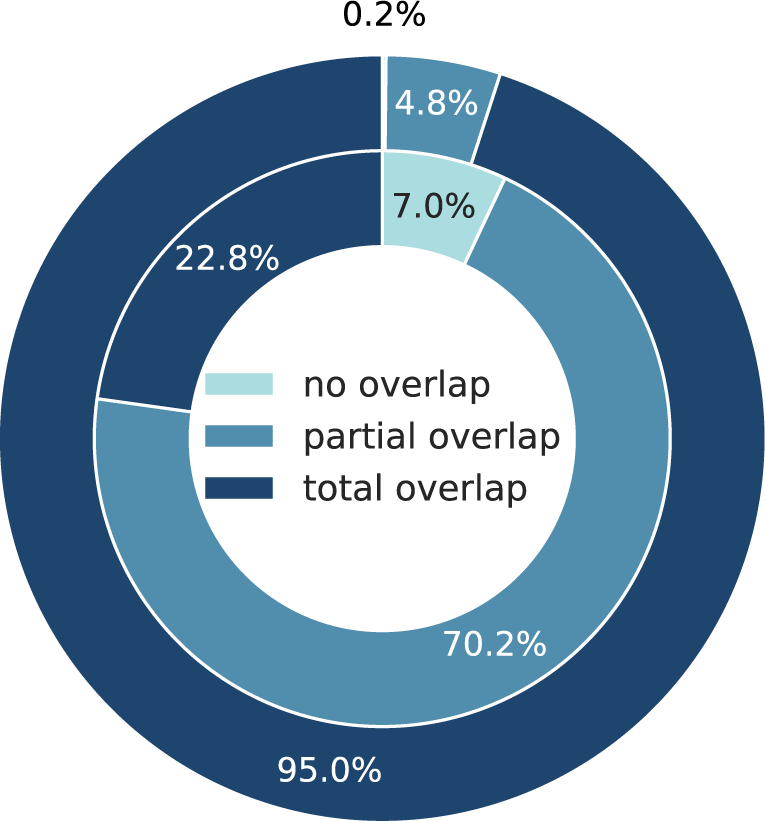
The impact of train/test splitting strategies on protein-level overlap. The common splitting strategy is to allocate pairs randomly (outer ring) while here we set aside proteins for testing (inner ring).

The choice of metrics is a crucial element of a benchmark, and summarising the results by a single number, such as accuracy or AUROC, is often misleading [33]. We report both the Receiver Operating Characteristic (ROC) and the Precision-Recall (PR) curves which highlight nuanced and complementary aspects of PPI models. In addition, to address the environmental impact of science in general [34] and bioinformatic tools in particular [35], we also reported the carbon footprint of training models, measured using the Green Algorithms calculator [36].

The elements described above represent the minimum needed for reproducible benchmarks and researchers who wish to use their own input features can evaluate their models on these partitions. However, to rapidly test a new model, it can be useful to have access to carefully curated protein properties. Amino acid sequences and functional genomics annotations, such as subcellular localisation and biological functions are available with B4PPI, from the professionally curated databases UniProt, the Human Protein Atlas (HPA) [37], [38] and Bgee [39] (Methods and Table 1 for the full list of features).

**Table 1:**
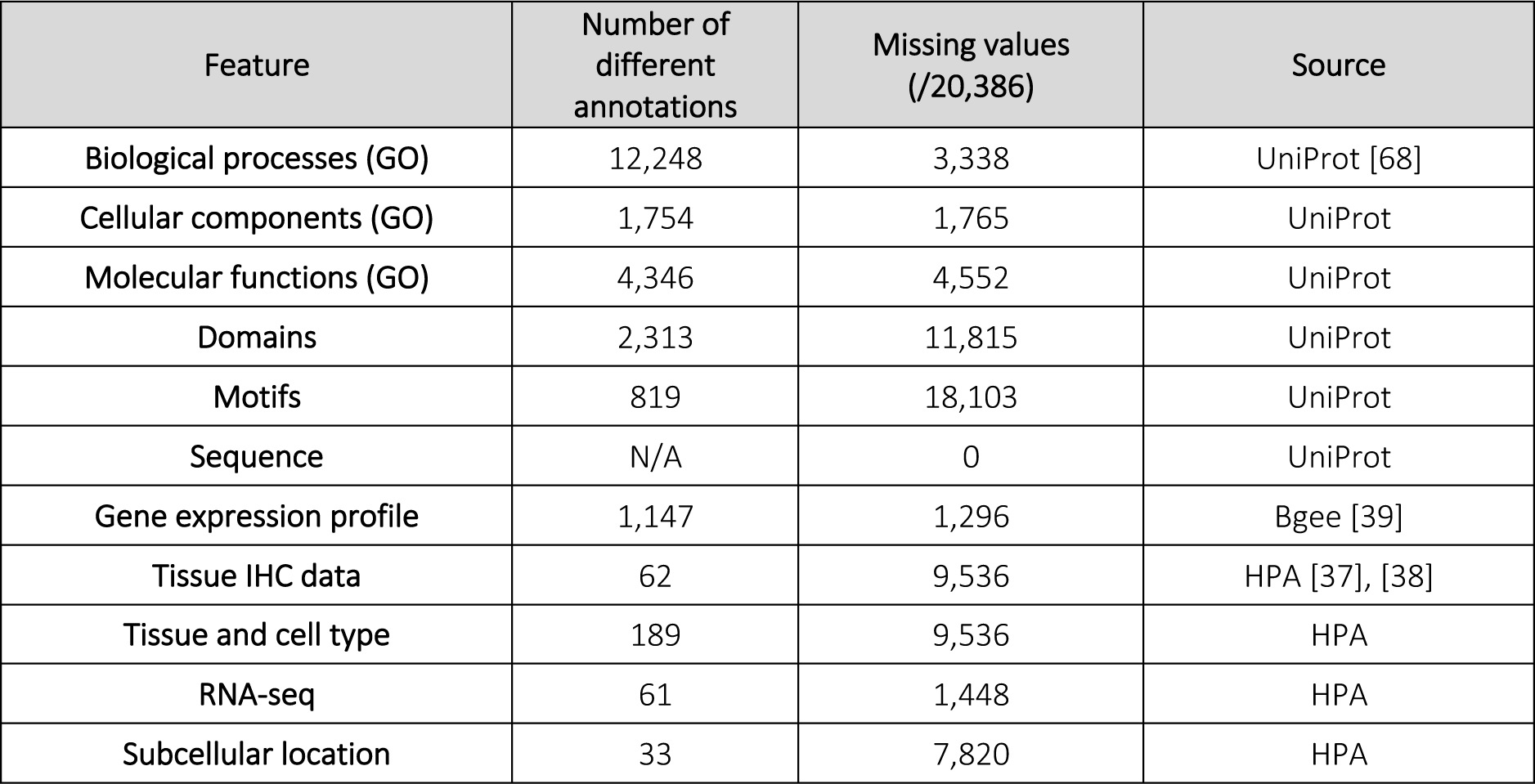
Features used to train human models (GO = Gene Ontology).

| Feature | Number of different annotations | Missing values (/20,386) | Source |
| --- | --- | --- | --- |
| Biological processes (GO) | 12,248 | 3,338 | UniProt [68] |
| Cellular components (GO) | 1,754 | 1,765 | UniProt |
| Molecular functions (GO) | 4,346 | 4,552 | UniProt |
| Domains | 2,313 | 11,815 | UniProt |
| Motifs | 819 | 18,103 | UniProt |
| Sequence | N/A | 0 | UniProt |
| Gene expression profile | 1,147 | 1,296 | Bgee [39] |
| Tissue IHC data | 62 | 9,536 | HPA [37], [38] |
| Tissue and cell type | 189 | 9,536 | HPA |
| RNA-seq | 61 | 1,448 | HPA |
| Subcellular location | 33 | 7,820 | HPA |

With B4PPI, we could compare different models in a consistent manner to better understand what aspects of the underlying biology are captured by each method. We focused here on FG-based and sequence-based models as they have been widely used and rarely compared, despite attempts at combining them.

### FG-based linear models achieved top performance

In FG-based models, FG annotations are pre-processed to compute similarity measures, such as colocalisation, between proteins (Methods and Table 1). The low dimensionality of the transformed problem explains the success of standard machine learning algorithms; in particular, Naïve Bayes Classifiers [40], decision trees [41] and Random Forests [42] have been the most popular choices [7]–[9], [43]. Despite the proven track record of such tools, the more recent XGBoost algorithm [44] has been shown to outperform them in other situations like kidney disease diagnostic [45], which motivated its inclusion in this analysis.

Using logistic regression as a baseline, we reported PR and ROC curves on the two test sets (Figure 1). A list of coordinates for these two curves was made available so that future models can be compared without unnecessary re-training. We also reported the training time, 6 seconds, and the carbon footprint, close to 0 gCO_2_e. We then compared other models to this baseline and produced similar performance sheets (Supplementary Figure 2).

We found that more complex algorithms brought little improvement over logistic regression, as most models performed similarly on *T1* (

Figure 3 and Supplementary Figure 3). XGBoost and Random Forest showed minor improvement in AUROC and AUPRC, but the difference between the ROC curves of the logistic regression and XGBoost is non-significant (p=0.27) (Methods). Moreover, XGBoost was more efficient than Random Forest as it had nearly half the runtime (30s vs 54s). When studying the coefficients of the linear regression, we found that most decisions are based on common biological processes, co-localisation (cellular compartment) and common domains, all three coefficients being significant (p<0.001) (Supplementary Figure 4). GO annotations for domains and motifs were missing for, respectively, 58% and 89% of proteins (Table 1), so we trained new FG-based models without them as a sanity check and found that it did not alter the results.

**Figure 3:**
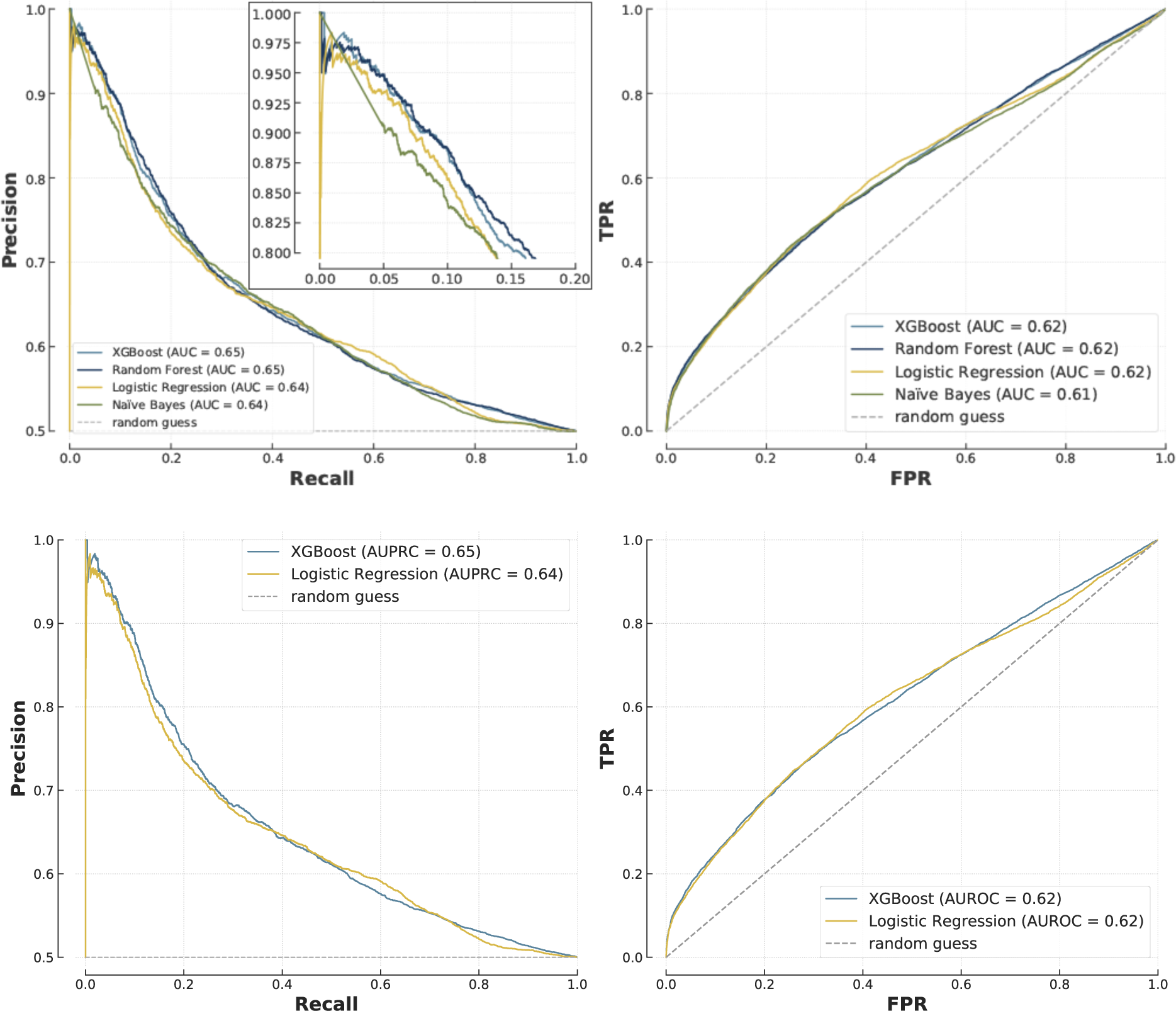
Comparison of FG-based models on *T1* (n=24,898, 50% positive), with PR curves (left) and ROC curves (right) for all the models tested (top) and then only XGBoost and Logistic Regression for clarity (bottom). The top left plot also shows a zoom on the high precision/low recall area.

The reporting standard also enabled us to look at finer performance metrics, broken down by protein-level overlap (i.e. individual proteins common to the training and testing sets). Comparing PR and ROC curves showed that both logistic regression and XGBoost were unaffected by the level of overlap (Figure 1 and Supplementary Figure 2), and can therefore transfer effectively to new proteins.

### Sequence models outperformed FG-based algorithms on known proteins

An alternative to FG-based models is to use amino acids sequences as the input for a PPI prediction algorithm. We compared several deep learning architectures and reported the performance of an optimised Siamese recurrent neural network, used in other recent PPI models [46] (Methods, Figure 9 and Supplementary Figure 5). Despite not having access to functional information about the proteins, the sequence model outperformed the best performing FG-based model, XGBoost, except at low recall and high precision (AUPRC=0.68 vs 0.65 and ROC curves significantly different, p=9x10^-47^) (Figure 4). However, while XGBoost was trained in only 30s with less than 0.01 kWh of energy, the deep learning approach trained for 1h10 with 0.62 kWh, emitting 22,000 times more greenhouse gases (GHGs). In addition, the performance of the sequence model was heavily affected by the choice of deep learning architecture and its hyper-parameters, such as number of layers or learning rate. These require extensive (and expensive) optimisation. We tested whether these results are impacted by sequence homology between proteins in the training and testing set (Methods), and we found that the sequence-based predictions on T1 are unaffected by sequence similarity (no correlation between prediction error and sequence similarity).

**Figure 4:**
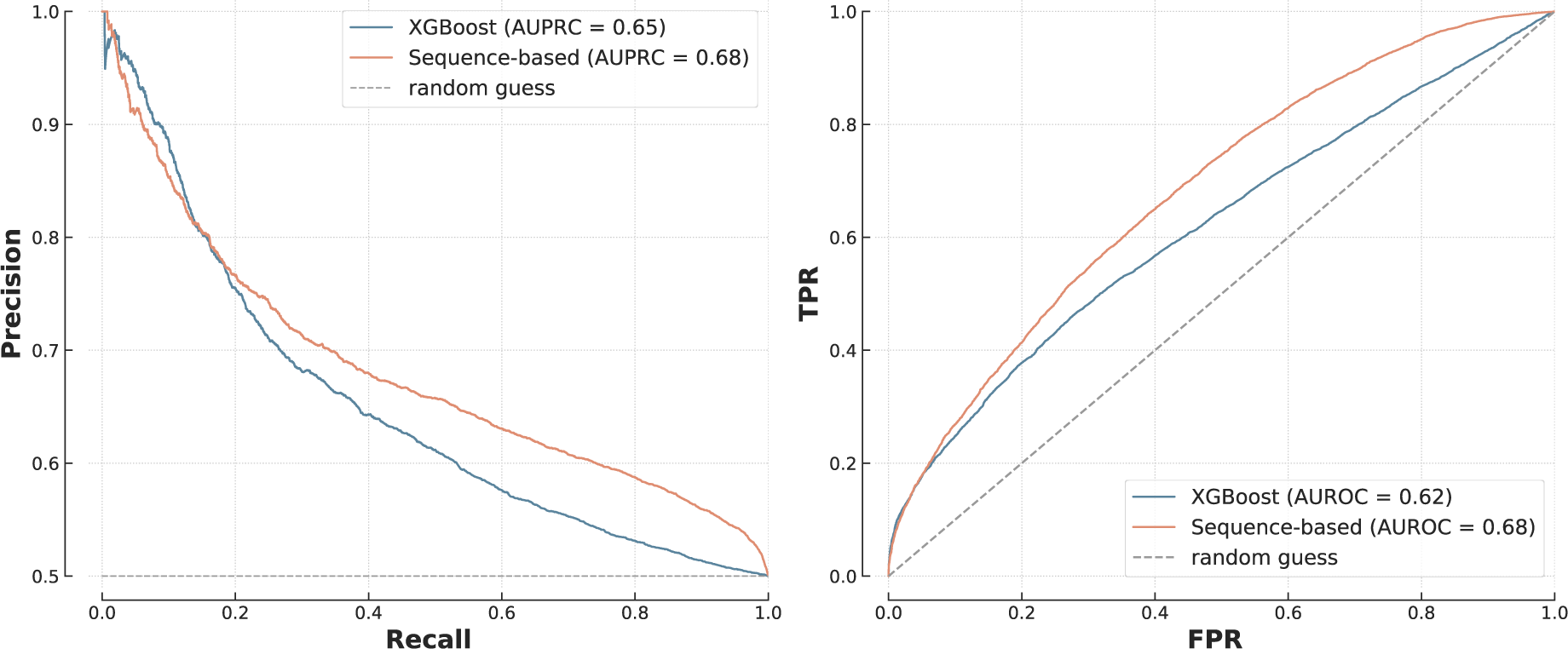
Siamese network vs XGBoost on *T1*. The difference between the ROC curves was statistically significant (p = 9x10^-47^).

Protein-level overlap had a significant impact on these results. The model had an AUPRC of 0.68 on average, but 0.75 when restricted to proteins present in both training and testing sets, and only 0.62 when there was no overlap (Supplementary Figure 5). This demonstrates that (1) in the absence of specific adjustments, such deep learning models are poorly suited to make predictions on previously unseen proteins and (2) in-depth benchmarks like B4PPI are important to reliably understand performance. While this comparison of FG-based and sequence-based models could indicate that deep learning is the best approach to PPI prediction, it could also be the consequence of unaccounted-for biological properties of PPIs.

### The role of network hubs is essential to PPI prediction

A scale-free topology has important biological implications [31] so we hypothesised that a one-fits-all approach for hubs and lone proteins is unlikely to be optimal. Contrary to network-based prediction models which are, by design, expected to be particularly sensitive to topology, the impact on sequence-based and FG-based models is not so evident. In assessing interactions between protein hubs (hub-hub), between a protein hub and a lone protein (hub-lone) and between lone proteins (lone-lone) (Methods), we found a distinct pattern whereby FG-based models had greater AUPRC and AUROC for interactions involving only lone proteins while sequence-based models performed better for hubs (Figure 5).

**Figure 5:**
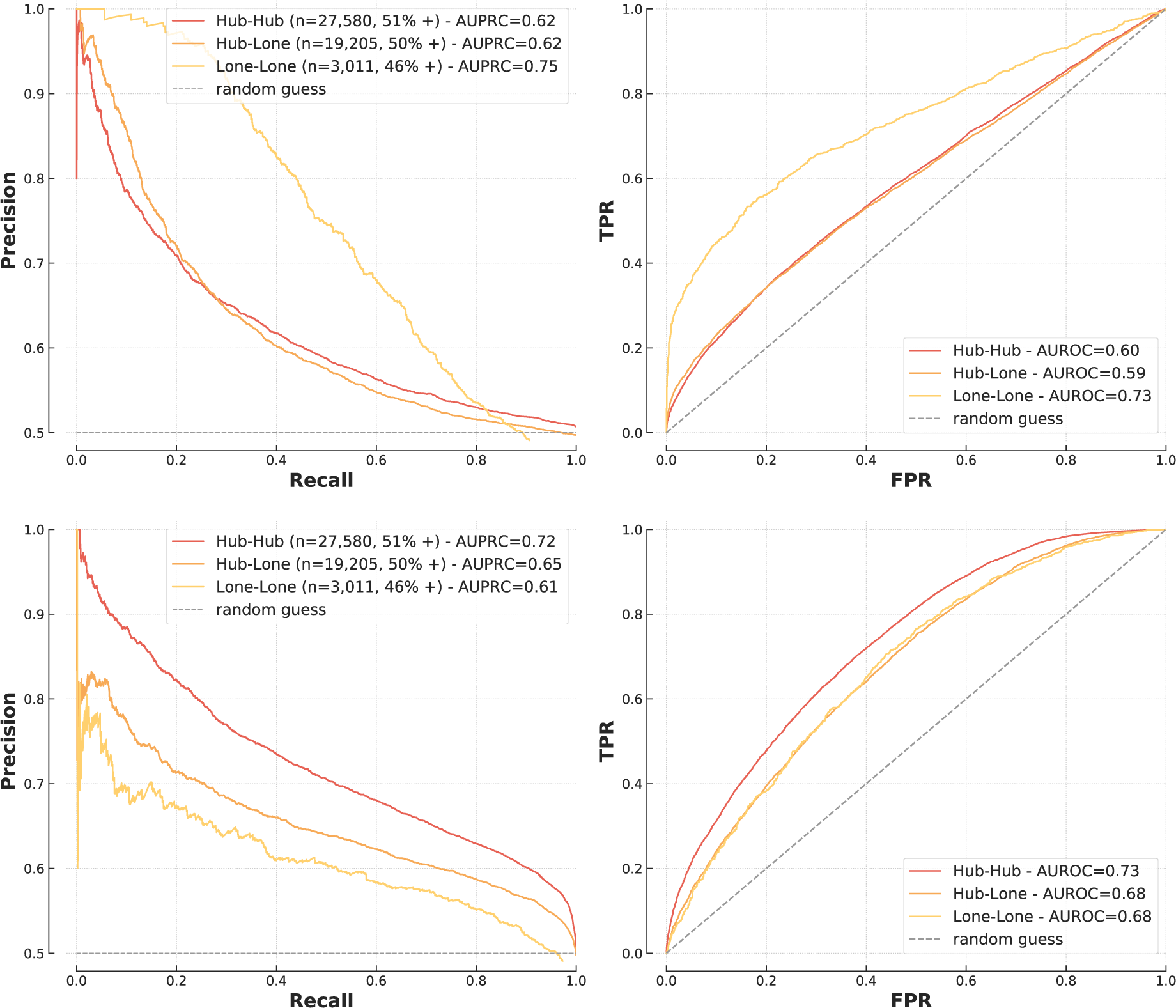
Performance of XGBoost (top) and sequence-based model (bottom) on hubs and lone proteins.

These findings can be explained by the pre-processing of similarity measures for FG models. Because of their central role in biological pathways, hubs are highly studied and therefore annotated for many processes and localisations. For example in the training set, hubs have on average 11.6 annotations for biological processes (significant feature in the logistic regression model discussed above) while non-hubs only have 5.8 (median 6 vs 3). The same phenomenon is observed for cellular compartments (6.2 vs 3.6 annotations on average). Because the similarity measures used by the FG models quantify overlaps in annotations, hubs annotated for a large number of processes provide little information about the probability of interaction, which can explain why FG-based models perform best when hubs are not involved.

These results provide insight into the strengths of each approach and, importantly, show that a PPI approach should be context specific, particularly with respect to the network topologies of interest. Indeed, the apparent superiority of the deep learning model shown on Figure 4 is largely due to the composition of *T1*, made up of 70% of hub-hub or hub-lone interactions.

### Cross-species validation of PPI prediction models and relative performances

*S. cerevisiae* is a well-studied model organism with a known interactome and has been used extensively for *in silico* PPI predictions [10], [43], [47]. We replicated the analyses presented above on *S. cerevisiae* proteins and found that our findings regarding network topology and models’ relative performances were robust across species. The data was selected similarly to previously, extracted and curated from IntAct and UniProt, but without data from HPA and Bgee as these databases do not curate yeast (Methods).

As shown previously, all FG-based models had similar performances with AUPRC between 0.71 and 0.73 (Figure 6); however, in this analysis, the differences between XGBoost and other models were statistically significant (p = 2x10^-12^ for Naïve Bayes and p = 9x10^-31^ for logistic regression). The sequence model outperformed FG-based models in most cases (p = 2x10^-6^), except at high recall (Figure 6). Second, similar to humans, FG-based models were not sensitive to protein-level overlap while sequence-based models had different performances depending on the level of overlap (Supplementary Figure 6). Finally, we found consistency regarding the role of network hubs; FG-based models were better able to predict lone-lone protein interactions while the sequence-based model was better at predicting interactions amongst protein hubs (Supplementary Figure 7).

**Figure 6:**
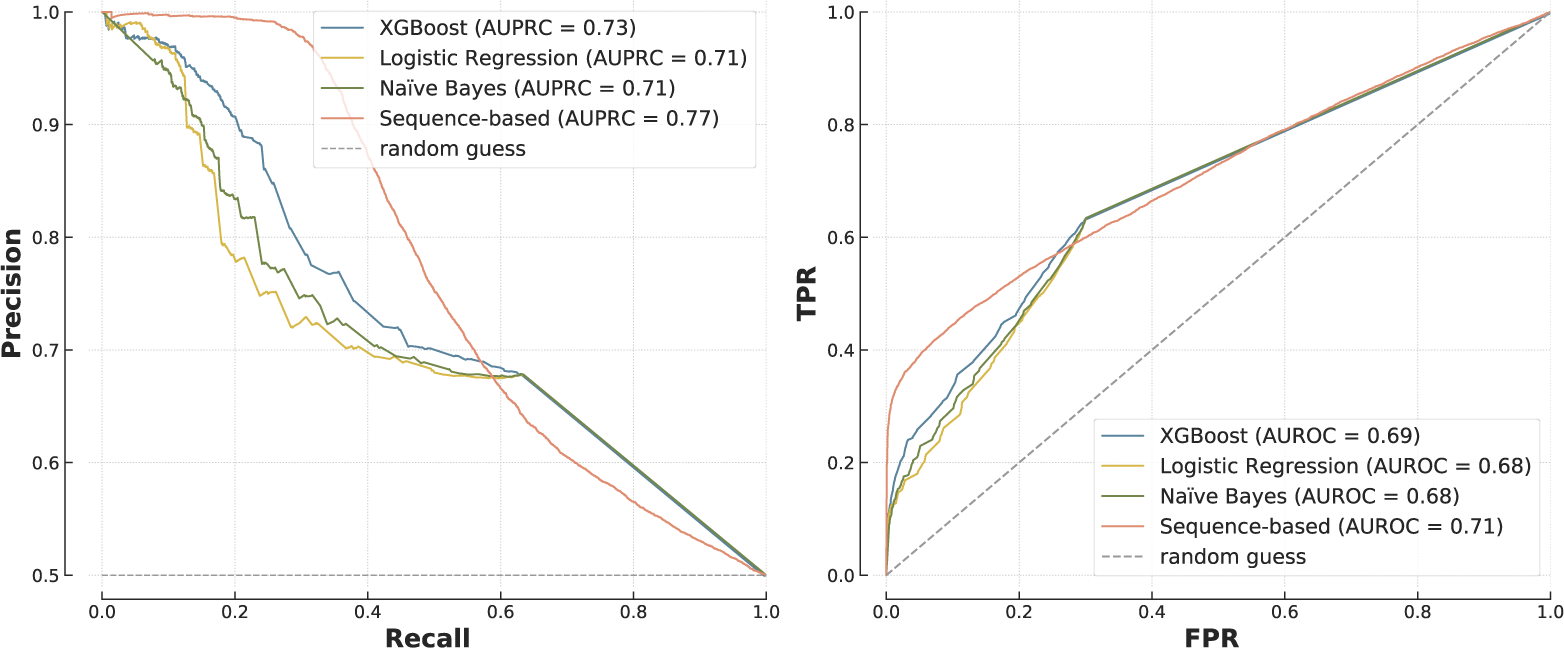
Comparison of a range of models on the yeast testing set.

While experimental data on PPIs is readily available for humans and *S. cerevisiae*, many non-model organisms lack data despite their biological relevance [48]. For these, cross-species predictions – i.e. training a model on a species to make predictions on another – are of particular interest. We showed that FG-based models are generally more suitable than sequence-based ones for this task.

We investigated whether models trained on yeast could be used to predict human PPIs, finding that the yeast-trained FG-based models (logistic regression and XGBoost) achieved similar AUPRC and AUROC as those which were human-trained to predict human PPIs (p = 0.26 for XGBoost) (Figure 7). Conversely, the yeast-trained sequence model was unable to predict human PPIs (AUPRC = 0.52 vs 0.68, p = 3x10^-272^). We observed the same phenomenon when using human-trained models to make predictions on yeast (Supplementary Figure 8).

**Figure 7:**
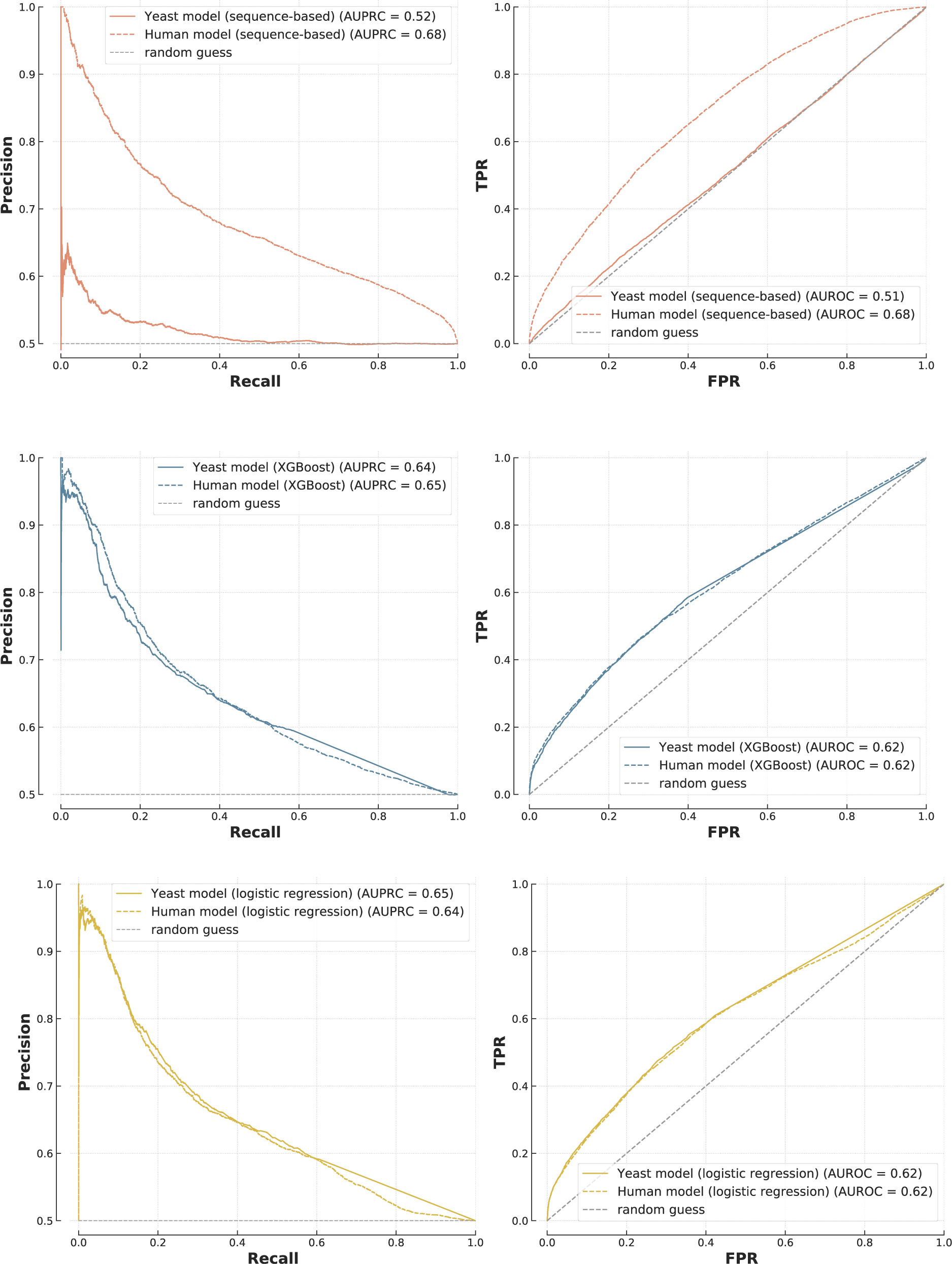
Cross-species predictions. Models trained on human PPIs (dotted lines) and yeast PPIs (solid lines) were used to make predictions on the human testing set. The top plot is the sequence-based model, the other ones are FG-based (XGBoost in the middle, logistic regression at the bottom).

## Discussion

In this work, we sought to identify and explain the strengths and weaknesses of PPI prediction approaches, and thereby provide the community with both a benchmarking pipeline and insight into which PPI approach to select or trust in a particular scenario.

The FG-based and sequence-based models are commonly used as building blocks for PPI prediction but are rarely directly compared. In particular, it was unclear where the differences lie and if one approach should be preferred today. The success of models based on gene ontology is in line with previous results [49], and we found that when using FG annotations, the choice of algorithm has little impact on the predictions and a logistic regression performs close to the state-of-the-art while providing clear insight into the decision-making process. Here, colocalisation, common biological processes and shared domains are the main indicators of interaction. Interestingly, different publications tend to highlight different FG features depending on the organism, dataset and predictor (mRNA expression in [8], coexpression/colocalisation in [50] or biological process in [49]). The fact that a highly flexible and non-linear model such as XGBoost performs similarly to logistic regression, making identical predictions in 93% of cases, shows that performance is likely driven by the quality and the pre-processing of the FG annotations instead of the modelling; once the similarity measures have been calculated, there are limited non-linearities and a simple logistic regression achieves top performance. Sequence-based models on the other hand need specifically optimised architectures but achieve similarly high performances, if not higher in some settings, without any biological information apart from amino acid sequences.

We found that the two approaches adapt to the presence of hubs and lone proteins differently and show complementary strengths. While sequence-based models are mostly useful when hubs are involved, FG-based models perform well for interactions between lone proteins. This simple result offers important insight into the specificities of each approach and explains discrepancies in reported performances in the literature, as the topology of the testing set has a large impact on metrics. These results are not specific to human PPIs as the same conclusions were drawn from analysis on *S. cerevisiae*. Cross-species predictions are instrumental to study non-model organisms, and we showed that FG-based decision rules translate well to new species while sequence-based models do not.

These observations are consistent with the way each algorithm learns. FG-based models make predictions based on general, but less complex, rules about PPIs which translate well to new proteins and new species. This is particularly useful considering that many proteins are still not represented in interaction databases. On the other hand, sequence-based models have millions of parameters which give them the flexibility to recognise individual proteins and learn specific interaction patterns. Although this enables such models to make predictions without functional information, it also limits high performance to proteins present in the training set, and make them particularly susceptible to data leakage [17]. This likely explains the poor results of sequence models on previously unseen proteins and cross-species datasets, something also observed by Dunham et al. in their benchmarking effort [16]. It is also consistent with the high performance of these models on network hubs, which are overrepresented in training sets and therefore well captured by the models.

These analysis and results required a robust and reliable benchmarking pipeline. We designed the open-source B4PPI, which accounts for a range of biological and statistical pitfalls. By being freely accessible and using standard identifiers for proteins, B4PPI can be used by any researcher working on *in silico* PPI prediction to investigate the inference mechanisms of their models. Contrary to previous benchmarking efforts [18], which it can complement, B4PPI leverages large professionally curated databases. An example reporting sheet is presented that includes relevant metrics, from PR and ROC curves to runtime and carbon footprint, to ensure the models released can be trusted and encourage wider use of PPI imputation for downstream analysis. B4PPI also comes with pre-processed features to enable rapid development of new approaches.

Our study has limitations. We focused on two widely used approaches to PPI prediction, namely FG-based and sequence-based; however, some alternative approaches have also been proposed, using, for example, higher-level protein structures [43], phylogeny [51] and the topology of existing networks [52]–[54], but the latter relies on the quality of the existing PPI networks. Most FG annotations are from gene ontologies which have a hierarchical structure which we do not account for here, contrary to Armean et al. [49] for example. Moreover, we analysed two common interactomes, human and yeast, yet there are many more. As demonstrated though with the yeast dataset, similar benchmarks and analysis can be transferred to other model organisms in a relatively straightforward manner. Literature-based databases such as IntAct are known to suffer from sampling bias as the curated studies tend to focus on already well-researched proteins [55], which may skew the predictions. One strategy to avoid sampling bias is to include high-throughput systematic mapping experiments, and IntAct - and therefore B4PPI - integrates a number of those, such as HuRI [5] and BioPlex [6].

We showed here the limits of classic sequence-based deep learning models on new proteins and for cross-species predictions, but it is worth noting some recent deep learning models that potentially addressed these limitations through careful regularisation for the former [46] and by including biological and chemical information about amino acids as well as structural knowledge for the latter [56], [57]. The results presented in this work can hopefully guide similar future work and help move this area further.

While benchmarking standards for PPI prediction are needed, it is important to remember the downsides of benchmarks, as demonstrated in computer vision or natural language processing. A fixed set of metrics can motivate the community to overly focus on those, at the expense of applicability and usefulness. To limit this, B4PPI includes a range of metrics but the relevant indicators for each use-case should nonetheless be carefully considered.

The size and complexity of the PPI network makes *in silico* prediction tools indispensable, but it is important to ensure that the models developed are reliable and readily available to the community for downstream analysis and to give insights into biological pathways. For this, consistent and reliable evaluation pipelines are necessary as well as a better understanding of what machine learning models learn. The results presented here make key progress in both areas and facilitate the development, evaluation and reliability of future PPI models.

## Methods

### B4PPI-Human

The data was obtained from large and professionally curated databases. This limits measurement bias, as each interaction is based on several experiments, and leverages experts’ knowledge in the curation process. On average, each PPI is supported by 3.4 publications (median of 3) and only 4.9% of interactions are obtained from only one source. Standard UniProt IDs are used throughout to ensure maximum compatibilities. Most of the manipulations were done in Python [58] with Jupyter Notebooks [59] using the Pandas library [60], [61] and Numpy [62]. The plots were drawn using Matplotlib [63], Seaborn [64] and the MetBrewer colour palettes [65]. All the code and final data are available on GitHub (https://github.com/Llannelongue/B4PPI); some intermediary pre-processed datasets are not available online due to file size limits but they can be recreated using the code available. Data is available under Creative Commons Attribution (CC BY 4.0) License.

#### Protein-protein interaction data

To train machine learning algorithms, the quality of the gold standard is paramount. Data on PPIs was obtained from IntAct [28] and downloaded from the EMBLE-EBI FTP server (timestamp: 15/10/2021). We restricted the data to human heterodimeric protein-protein interactions with UniProt IDs. To reduce the risk of false positives, we removed complex expansions (where the pairwise interactions within a complex are unreliable) and interactions based on colocalisation only. This quality control step leaves 128,790 PPIs, covering 15,506 proteins (out of 20,386 in UniProt). Based on this dataset, we created an index of the number of recorded interactions per protein and made a list of hubs (highly connected proteins). In line with the literature, hubs are defined as the 20% of proteins with the most interactions [66], which here is equivalent to proteins with more than 21 partners. The quality of the interactions is assessed further by looking at the MIscore [67], a quality score based on the manual curation of the interactions and annotations of the HUPO PSI-MI consortium that takes into account the detection method, the interaction type and the number of publications reporting it. In case of PPIs with multiple entries, the highest MIscore was used. When looking at the distribution of the MIscores in the dataset (Figure 8), a threshold of 0.47 is visible, which restrict the dataset to 78,229 interactions, covering 12,026 proteins. We also ran the same analyses without filtering on MIscores (i.e. using all 128,790 interactions) and found that all the results presented here held true.

**Figure 8:**
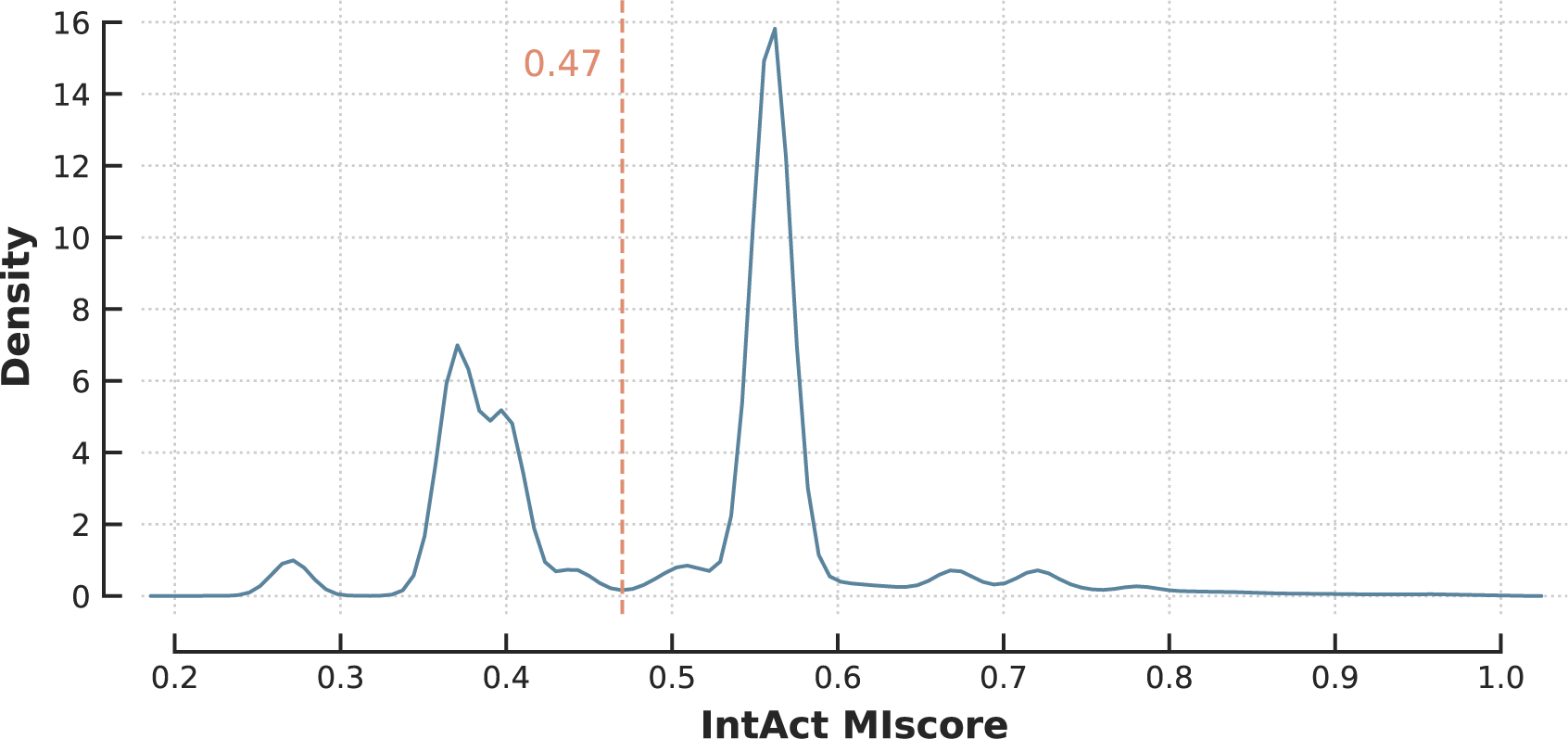
Distribution of the MIscore in IntAct.

#### Functional genomics annotations and amino acids sequences

Protein sequences in humans are well documented and can be obtained from UniProt, but FG features can be more challenging as they should be diverse (i.e. cover a wide range of properties), of high-quality and have high coverage (i.e. few missing proteins). For the same reasons as described above, aggregated, manually curated, and professionally reviewed databases are preferred. Based on features that have been successfully used for the task before, it is relevant to include information about cellular and tissue localisation, biological functions and gene expression patterns [7], [10], [12], [43].

One of the main databases on proteins is UniProt [68] and in particular its knowledgebase UniProtKB. Swiss-Prot, the section of UniProtKB that is reviewed and manually curated, is used in this work to ensure optimal quality. The data from Swiss-Prot is downloaded through their API by restricting to reviewed, non-obsolete, human proteins (last download is 09/11/2021). The different columns are then cleaned to extract the information of interest in a standardised format, and we use UniProt IDs throughout. There is information for 20,386 proteins and more details about each feature are in Table 1. UniProt’s API is also used to map UniProt IDs between different databases and to map outdated IDs. In particular, we extract amino acids sequences for each protein, more than 95% of which come from the translation of coding sequences submitted to the International Nucleotide Sequence Database Collaboration [69]. Annotated domains and motifs are also included in the database. Additionally, we extract gene ontology (GO) annotations of biological processes, cellular components and molecular functions. For each protein, each of the FG features is represented as a bag-of-words, i.e. a sparse vector of length the number of annotations in the database.

When working with gene expression data, both biological and technical noise need to be accounted for correctly. The Bgee public repository [39] does that by regrouping curated healthy wild-type standardised gene expression patterns. The human data is mainly from GTEx v6 (phs000424.v6.p1), with an added layer of manual curation to remove unhealthy subjects. For a gene, the final data provides binary calls of presence or absence of expression for each combination of anatomical entity and developmental stage. We downloaded the database from their FTP server (version 14.2) and obtained information for 59,777 genes, 320 anatomical entities and 33 developmental stages, which leads to 1,147 stage/entity combinations. The Bgee entries are matched to the UniProt IDs using UniProt’s own mapping table.

The Human Protein Atlas (HPA) [37], [38] provides data mapping human proteins to tissues and cells. In particular, we used the Tissue Atlas [37] that presents the distribution of proteins in tissues and cell types and the Cell Atlas [38] that contains the distribution across subcellular locations. The Tissue Atlas contains data similar to Bgee, but the overlap is likely to be limited as the two databases only share GTEx RNA-seq data. While Bgee has a more thorough curation process, HPA contains a lot of original in-house experimental results, which justifies the inclusion of both data sources. We downloaded the HPA data from their website (release 20.1, Ensembl version 92.38). The data in HPA is identified by Ensembl gene IDs, which are mapped to UniProt IDs using UniProt’s API. We restricted the dataset to the reviewed proteins present in Swiss-Prot and to ensure the quality of annotations, we discarded the entries HPA annotated as “uncertain”. For the tissue IHC data, we mapped expression levels to numerical values (high=3, medium=2, low=1 and “not detected”=-1) with untested tissues being mapped to 0. Similar pre-processing was used for the consensus RNA-seq data and the subcellular location.

#### Pre-processing to measure features similarity

For a protein, each FG feature was represented as a vector, of length the number of annotations. To measure the feature-specific similarity between two proteins, we compared the two vectors using cosine similarity [70], a popular tool widely used for similar tasks in Natural Language Processing. For two vectors *A* = (*A_i_*) and *B* = (*B*_i_), their cosine similarity *CS*(*A*, *B*) is:

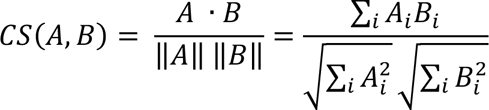

As a result, for each of the 207,784,305 possible pairs of proteins, we obtained 12 similarity features: biological processes, cell components, molecular function, domains and motifs from UniProt, gene expression from Bgee, tissue/cell expression, tissue expression, RNA-seq expression and subcellular locations from the Human Protein Atlas (Table 1).

#### Creation of the gold standard

The PPIs obtained from IntAct are divided between a training set and two testing sets. First, a set of 1,562 proteins (13%) was randomly set aside to ensure some unseen proteins are present in the testing set; the necessity of this is shown in Figure 2. This percentage was chosen as it gives a train/test ratio of ∼70%/30%, which is standard when data is limited. The dataset was then randomly divided under this constraint and included 53,331 PPIs in the training set, 12,449 in *T1* and the same in *T2* (Supplementary Table 1). We tested the sensitivity of the results to the division parameters by creating new gold standards where 10% and 15% of proteins were set aside for testing (instead of the 13% mentioned above) and found that all the results remained identical.

The negative examples (i.e. non interacting proteins) are obtained using random sampling among all the possible pairs, excluding any pair that has been observed experimentally to limit the risk of false negative. For the training set, balanced sampling is used [32] to favour learning, which means that the probability of sampling a protein for the negative set is proportionate to its frequency in the positive set. For *T1* and *T2*, we used uniform sampling (all proteins have the same probability of sampling) to limit sampling bias. The training set and *T1* both have 50% of positive examples, while *T2* has only 1% of interacting pairs (Supplementary Table 1).

We looked at sequence homology between proteins in the training and testing sets to ensure sufficient independence, using Ensembl’s paralogs database. We found that 47% of *T1* proteins have no paralogs in the training set and among the remaining 53%, the average similarity between two proteins is 25%. Besides, 70% of paralogs pairs have less than 30% similarity. This indicates that the testing set is sufficiently independent.

To investigate how models deal with different network topologies, especially hubs and lone proteins, we had to create a separate testing set to ensure sufficient sample size in each category (hub-hub, hub-lone and lone-lone interactions). We do so by aggregating PPIs from *T1* and *T2*, and using balanced sampling for the non-interacting proteins. This results in 49,796 pairs (50% positive) (Supplementary Table 2).

### S. cerevisiae data

The pipeline describe above was also followed for the *S. cerevisiae* data. UniProt lists 6,721 yeast proteins and the same information as for humans (Supplementary Table 3) but HPA and Bgee do not include data for this organism. PPIs were obtained from IntAct following the same procedure, although no selection based on MIscores was made considering the absence of an obvious choice when looking at the distribution (Supplementary Figure 9). The final PPI dataset comprised 43,068 interactions covering 5,679 proteins.

The split between training and testing sets was done similarly by setting aside 737 proteins for testing and then randomly allocating PPIs to keep 30,369 PPIs for training (70% of the gold standard). Because there is fewer data on yeast, and only one testing set is needed to replicate the analysis conducted on humans, dividing the remaining 12,699 further between *T1* and *T2* is not suitable here. But if the goal was to measure generalisability of a yeast model, this could be easily done.

### Training

FG-based machine learning models were trained using the scikit-learn library [71]. For models that cannot deal with missing data, mean imputation was used (Supplementary Table 4). Hyper-parameter search was done using Weight-and-Bias’s Bayesian method [72] to find the optimal settings of each algorithm in a reasonable time. All hyperparameter choices are in Supplementary Table 4 and Supplementary Table 5.

Deep learning models were trained using PyTorch Lightning [73], [74]. The Siamese architecture [75], [76] was composed of a bidirectional Gated Recurrent Unit (GRU) [77] followed by a linear output (Figure 9). Long-Short Term Memory networks (LSTM) [78] and Convolutional Neural Networks (CNN) [79] were also tested, but GRU was preferred because of runtime efficiency and its ability to account for proteins of various lengths. Full parameters are in Supplementary Table 4, Supplementary Table 5 and in the open-source code.

**Figure 9:**
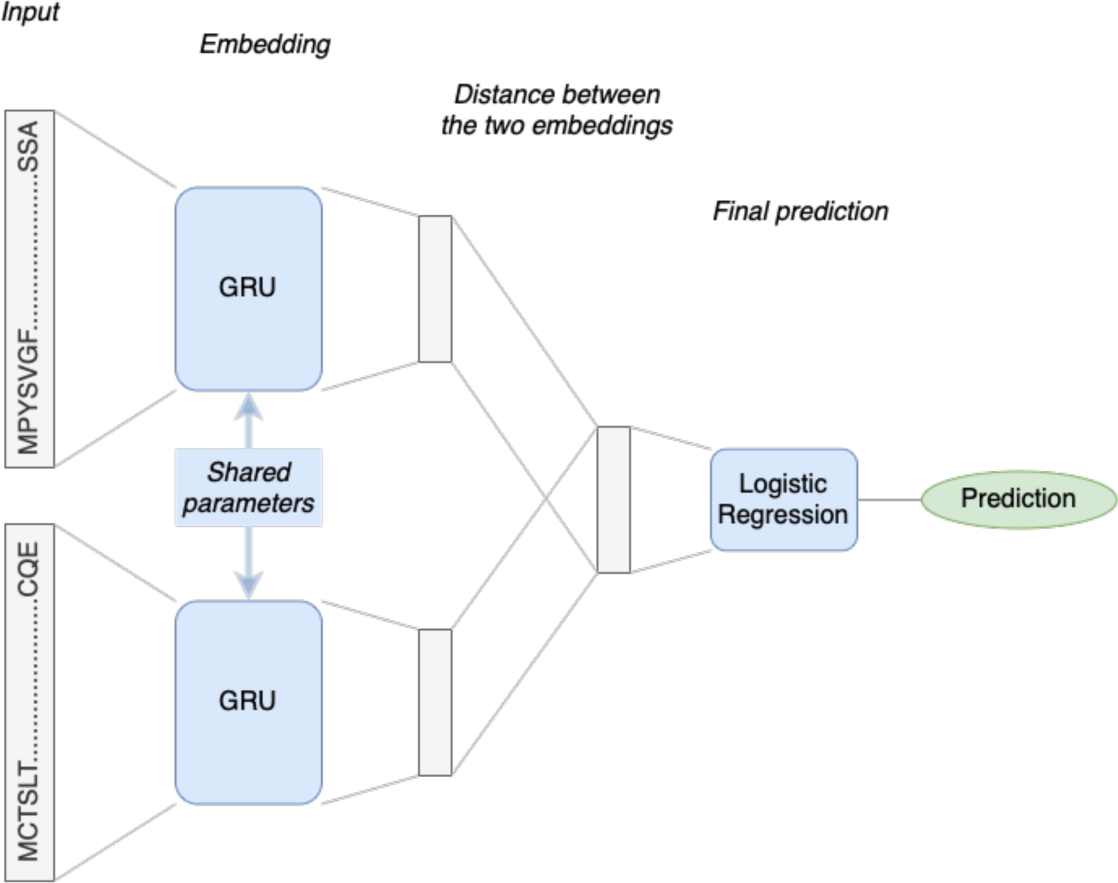
Diagram of the deep learning architecture used to predict interactions from a pair of protein sequences.

### Evaluation

The Receiver Operating Characteristic (ROC) and the Precision-Recall (PR) curves are complementary options for PPI prediction. While the ROC curve is unaffected by the prevalence of interacting proteins, a benefit as the true prevalence of PPIs is mostly unknown, it also means that both classes are considered equally, whereas often, PPIs are more interesting than non-interacting proteins. This is addressed by the PR curve where precision puts an emphasis on positive examples. It has also been shown that ROC tend to overestimate the performance of PPI prediction tools [18], but most published models still report it, which justifies the inclusion of both metrics.

Both curves are reported alongside their respective Areas Under the Curve (AUC). To statistically compare ROC curves for a same testing set, we used a DeLong nonparametric test [80] and reported the p-value. We corrected for multiple testing by using a conservative significance threshold of 5 x 10^-4^, corresponding to a Bonferroni correction for 100 pairwise comparisons [81].

### Environmental impact statement

We did our best to minimise greenhouse gas emissions related to this project and, using the Green Algorithms calculator (v2.1) [36], we estimated that the carbon footprint of this project was 51 kgCO_2_e, which corresponds to 4.7 tree-years.

## Acknowledgments

L.L. was supported by the University of Cambridge MRC DTP (MR/S502443/1) and the BHF programme grant (RG/18/13/33946). M.I. was supported by the Munz Chair of Cardiovascular Prediction and Prevention and the NIHR Cambridge Biomedical Research Centre (BRC-1215-20014) [*]. MI was also supported by the UK Economic and Social Research 878 Council (ES/T013192/1).” This work was supported by core funding from the British Heart Foundation (RG/13/13/30194; RG/18/13/33946) and the NIHR Cambridge Biomedical Research Centre (BRC-1215-20014) [*]. *The views expressed are those of the author(s) and not necessarily those of the NIHR or the Department of Health and Social Care.

This work was also supported by Health Data Research UK, which is funded by the UK Medical Research Council, Engineering and Physical Sciences Research Council, Economic and Social Research Council, Department of Health and Social Care (England), Chief Scientist Office of the Scottish Government Health and Social Care Directorates, Health and Social Care Research and Development Division (Welsh Government), Public Health Agency (Northern Ireland), and British Heart Foundation and Wellcome.

For the purpose of open access, the author has applied a Creative Commons Attribution (CC BY) licence to any Author Accepted Manuscript version arising from this submission.

## Supplementary material

### Supplementary Figures

**Supplementary Figure 1:**
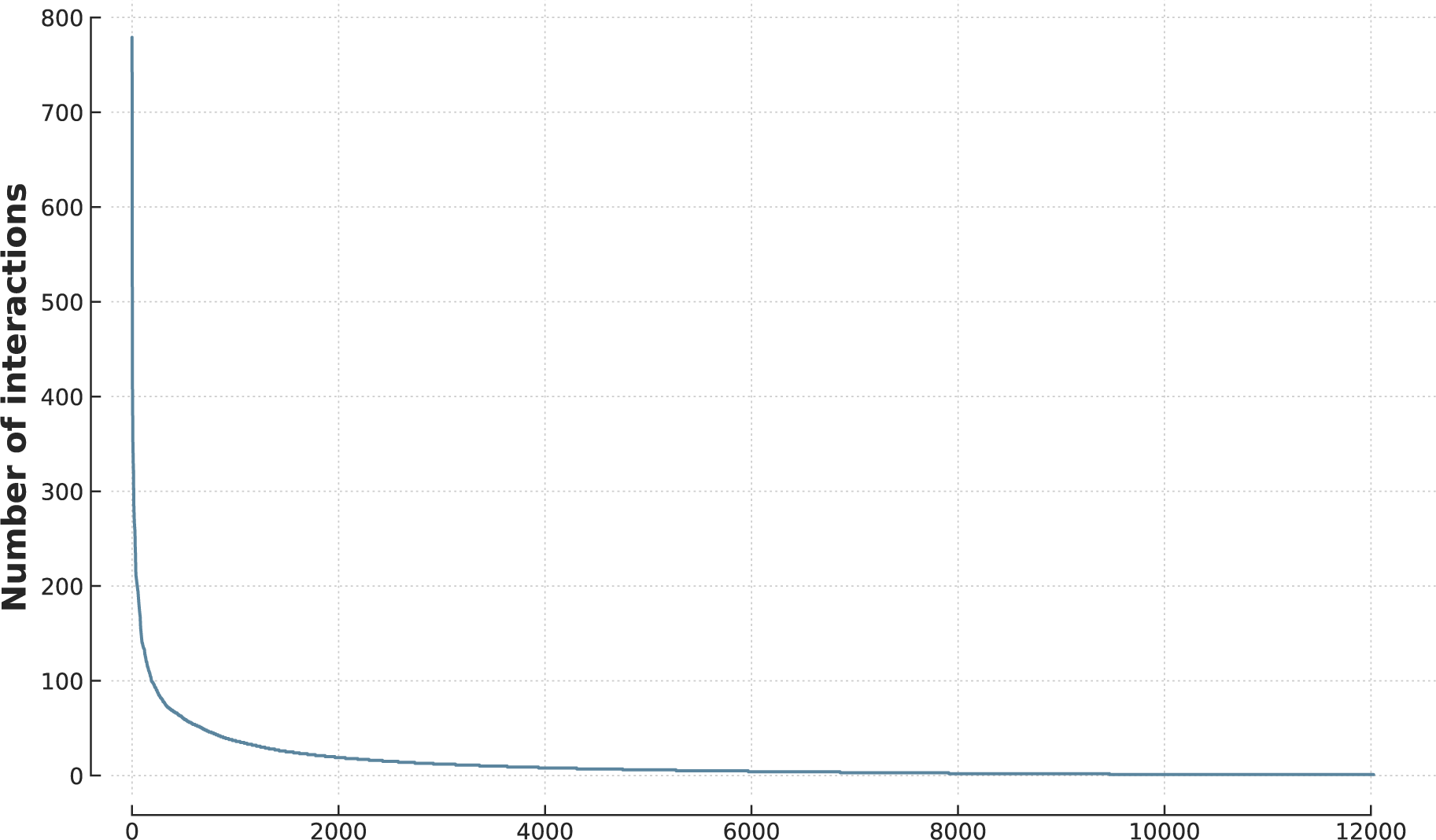
Distribution of proteins’ degree in IntAct. The exponential decrease is characteristic of a scale-free network.

**Supplementary Figure 2:**
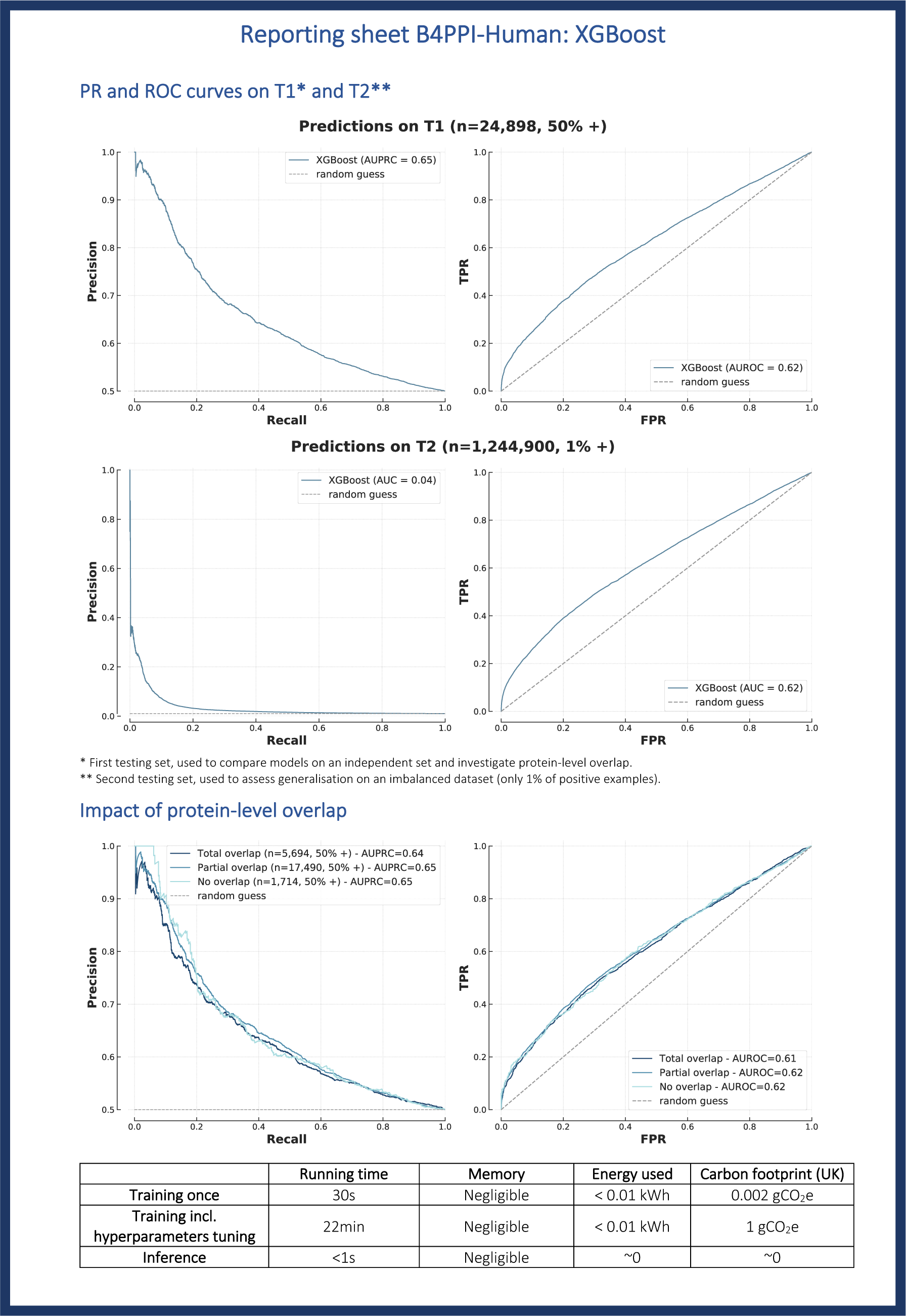
Performance sheet of XGBoost on B4PPI-Human.

**Supplementary Figure 3:**
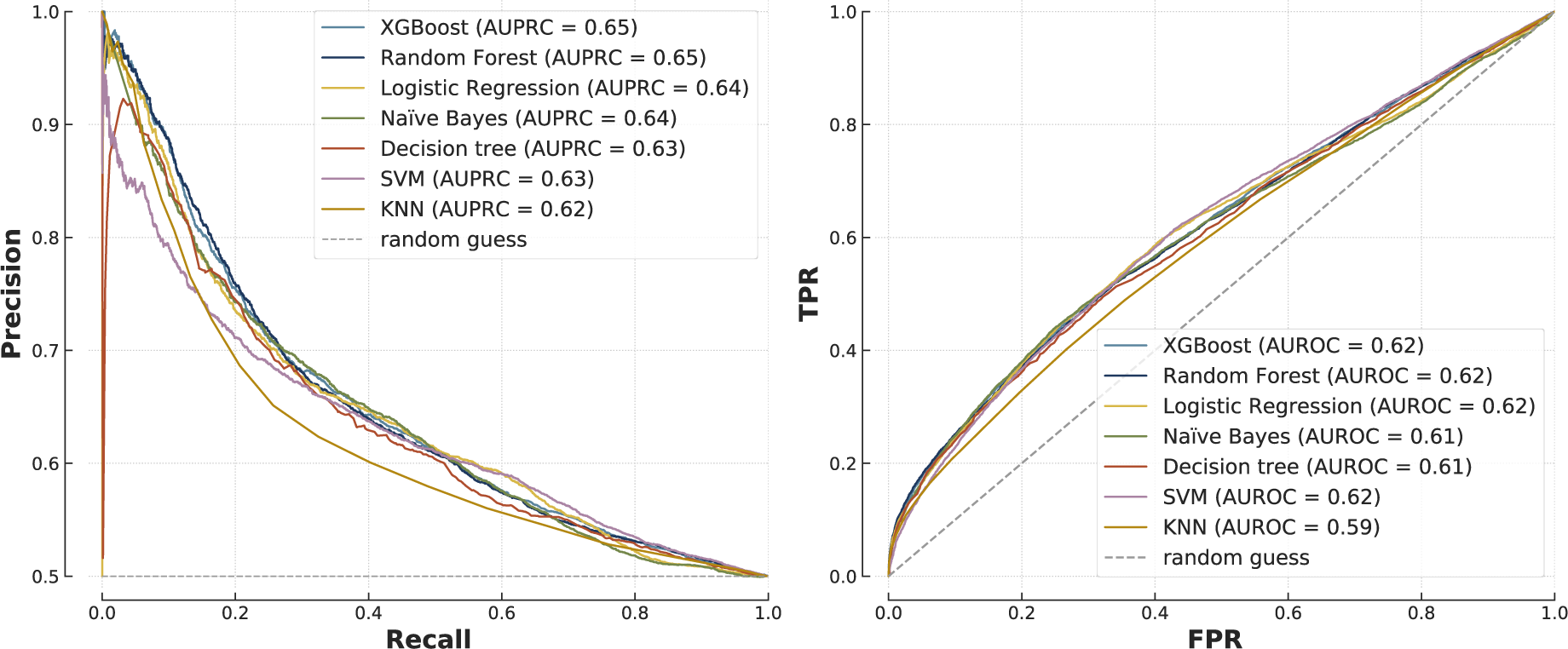
Comparison of a broader range of FG-based models on T1.

**Supplementary Figure 4:**
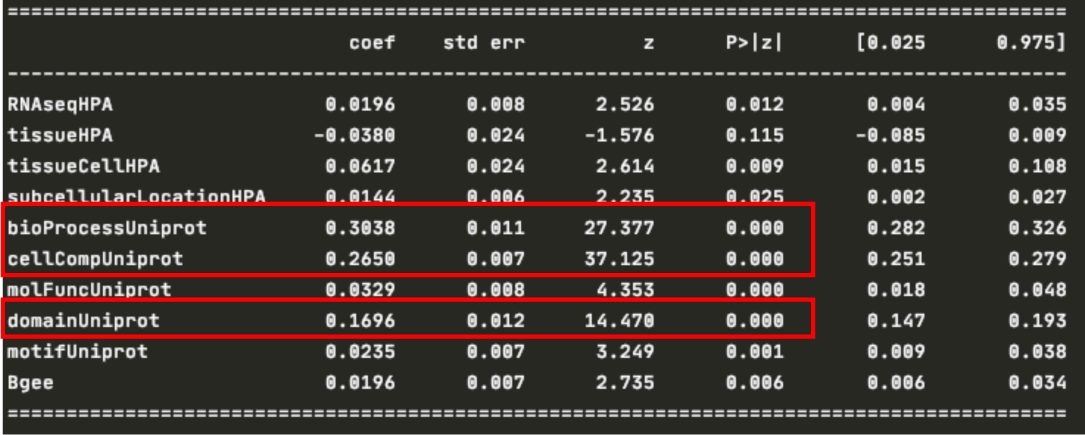
Output of the logistic regression on the training set.

**Supplementary Figure 5:**
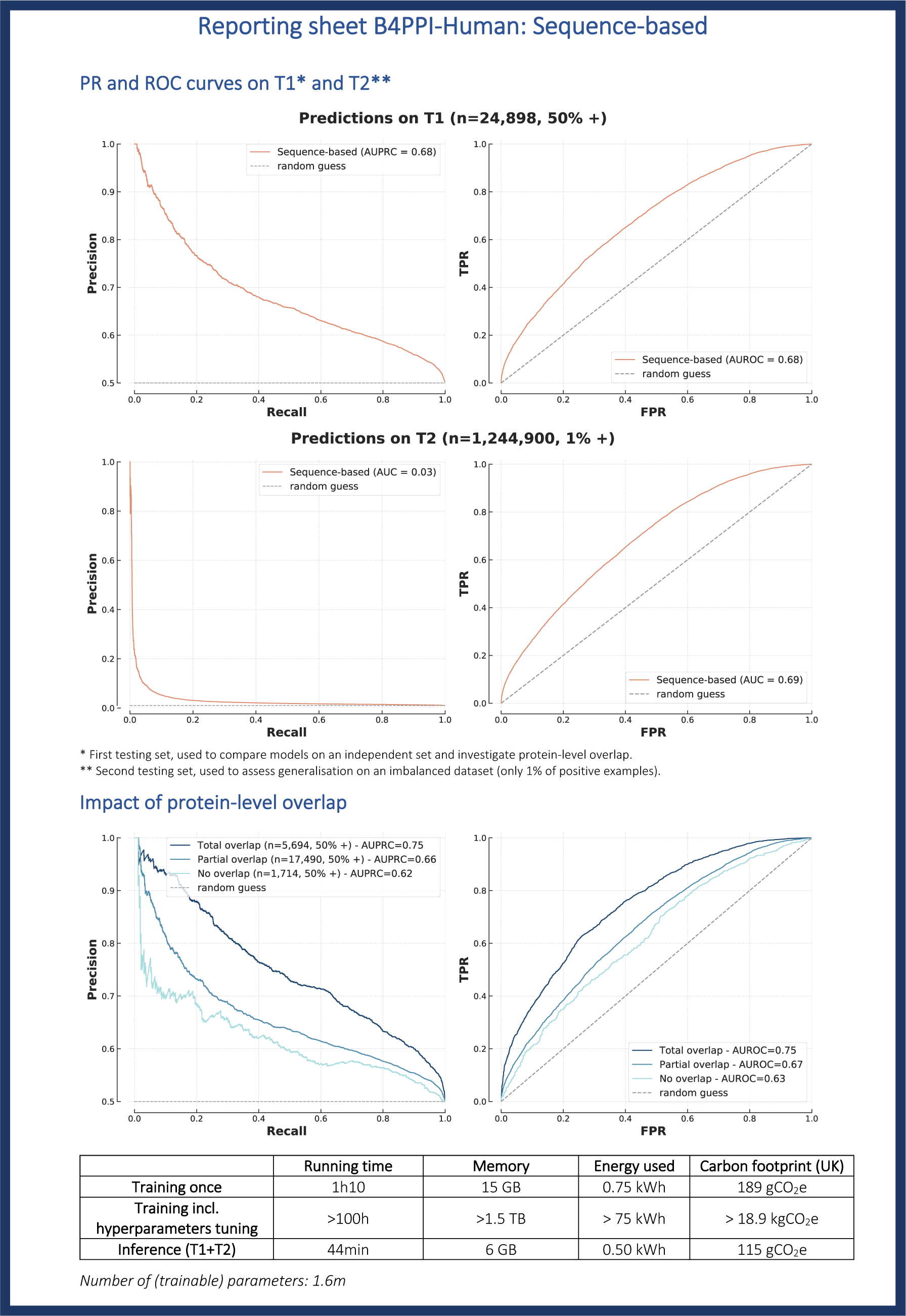
Performance sheet of the sequence-based model.

**Supplementary Figure 6:**
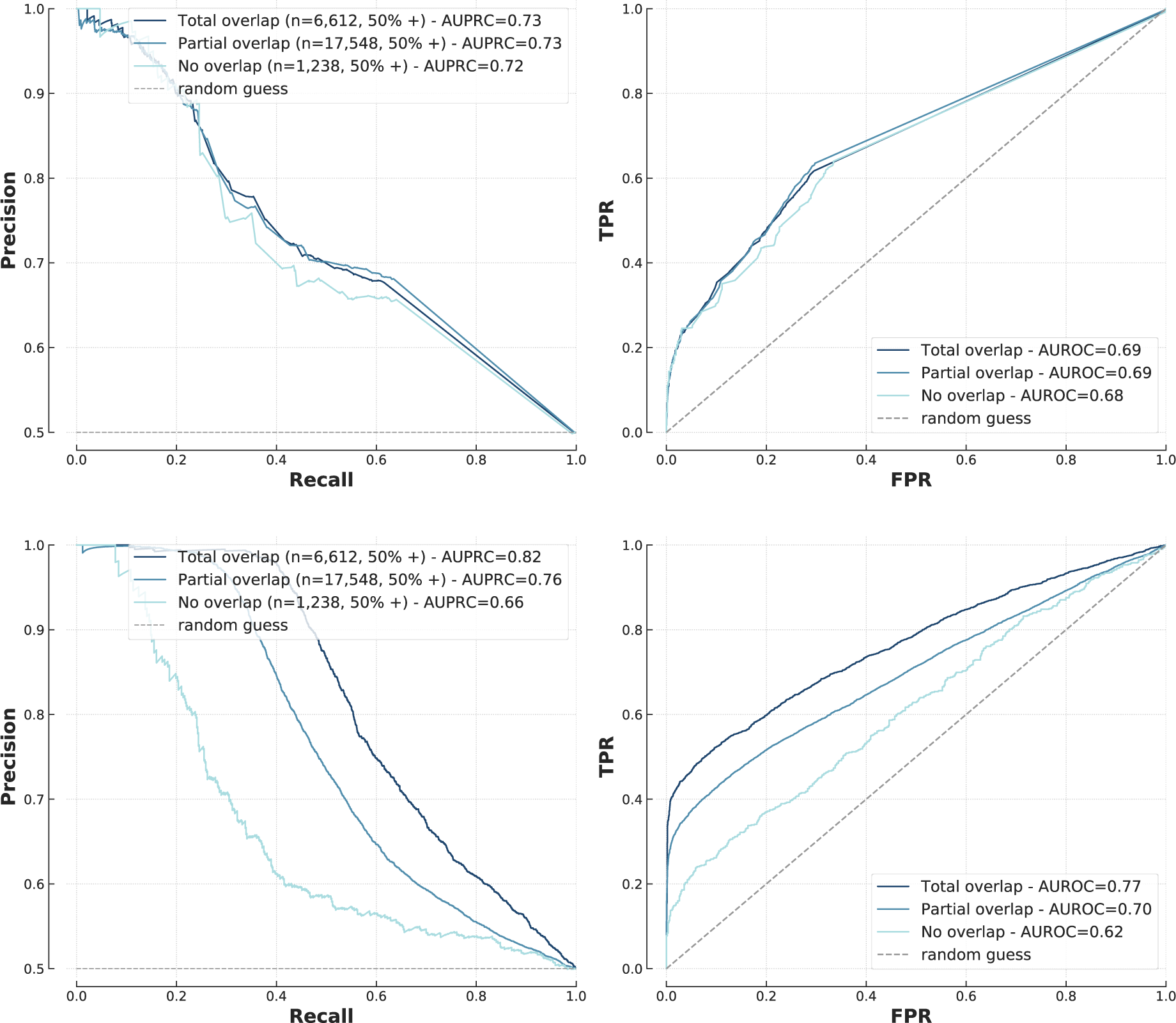
Impact of protein-level overlap on the yeast dataset for XGBoost (top, FG-based) and the sequence-based model (bottom).

**Supplementary Figure 7:**
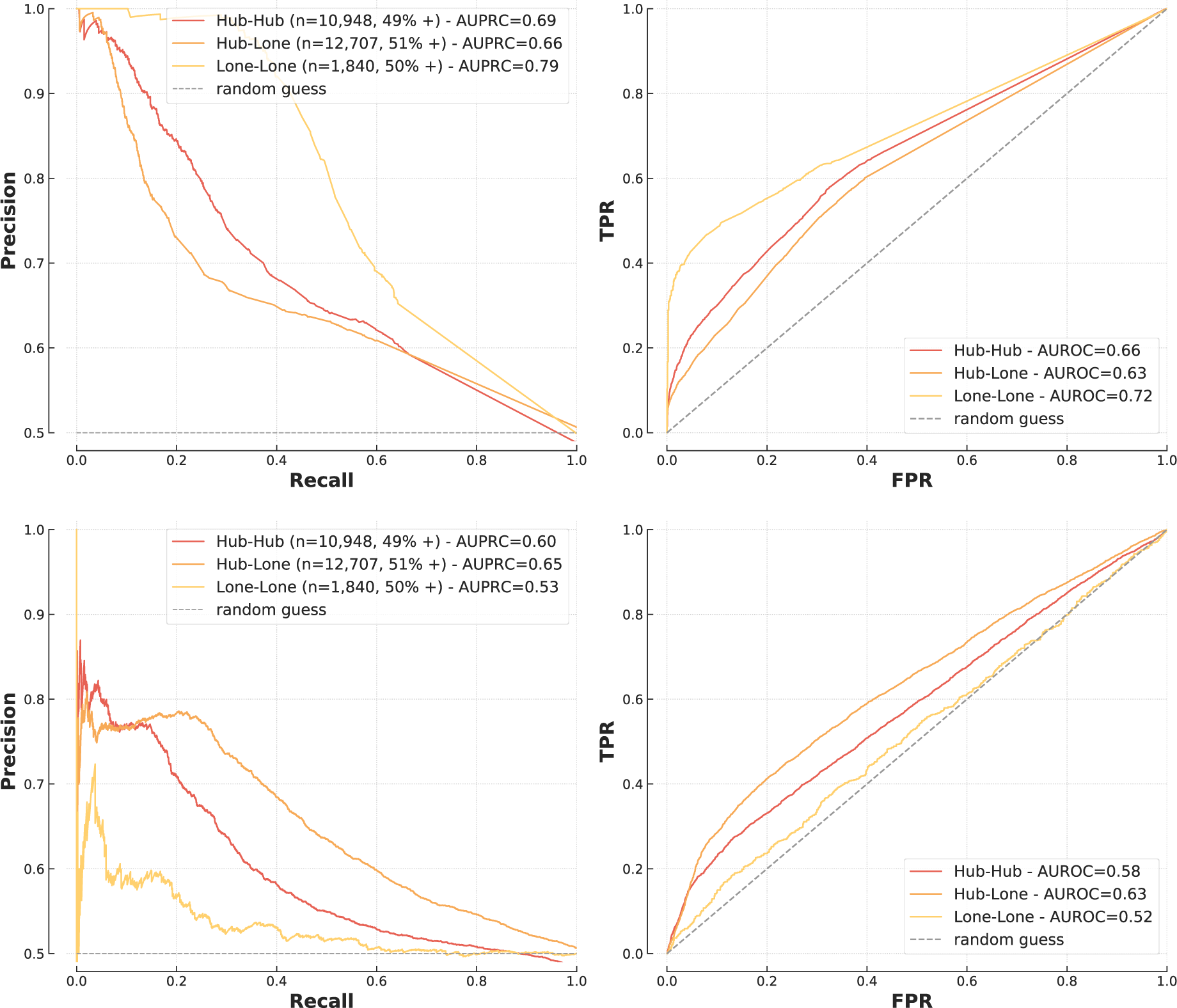
Impact of hubs for FG-based (XGBoost, top) and sequence-based (bottom) model on yeast interactions.

**Supplementary Figure 8:**
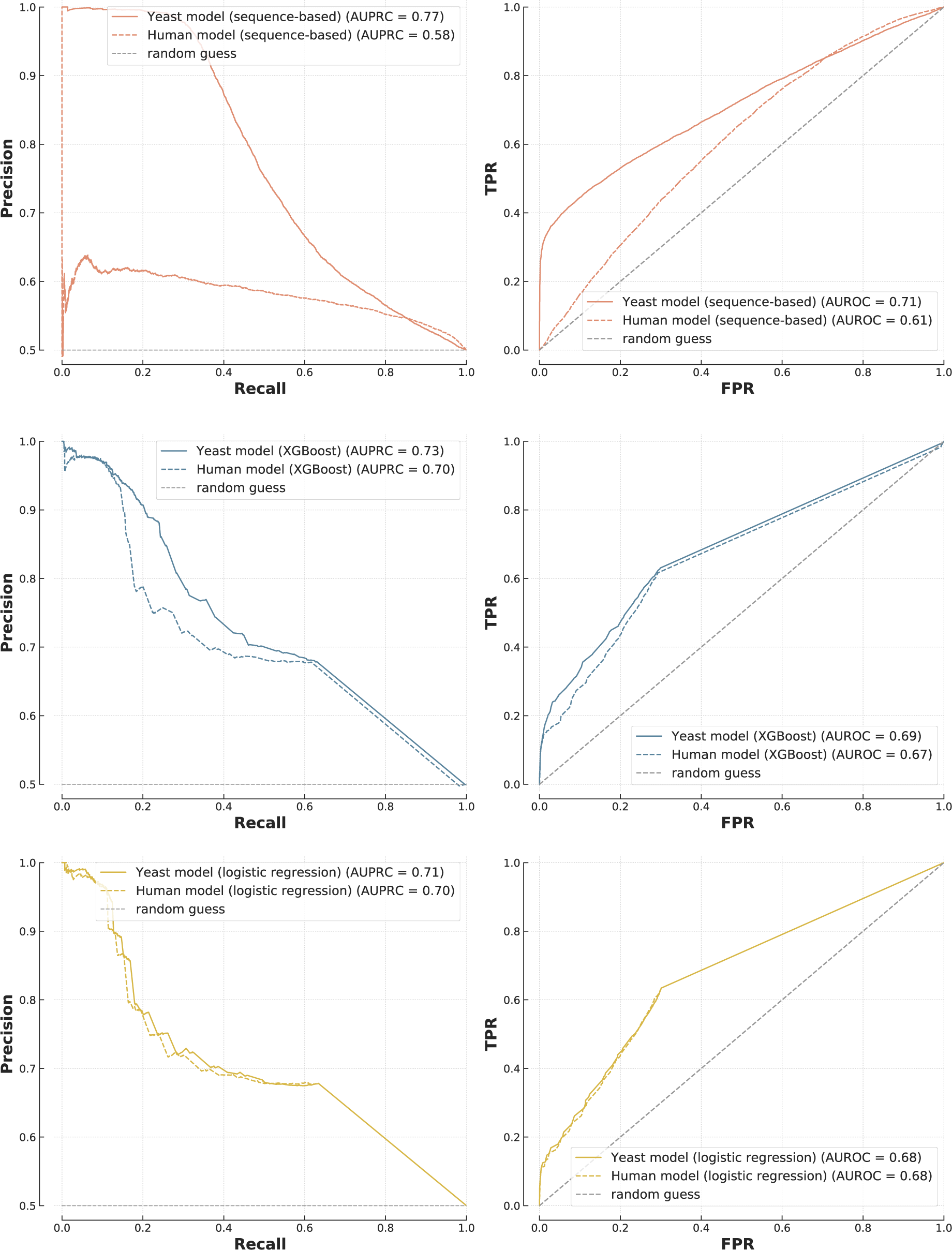
Cross-species predictions. Models trained on human PPIs (dotted lines) and yeast PPIs (solid lines) were used to make predictions on the yeast testing set (n=25,398, 50% positive). The top plot is the sequence-based method, the others the FG-based ones (XGBoost in the middle, logistic regression at the bottom).

**Supplementary Figure 9:**
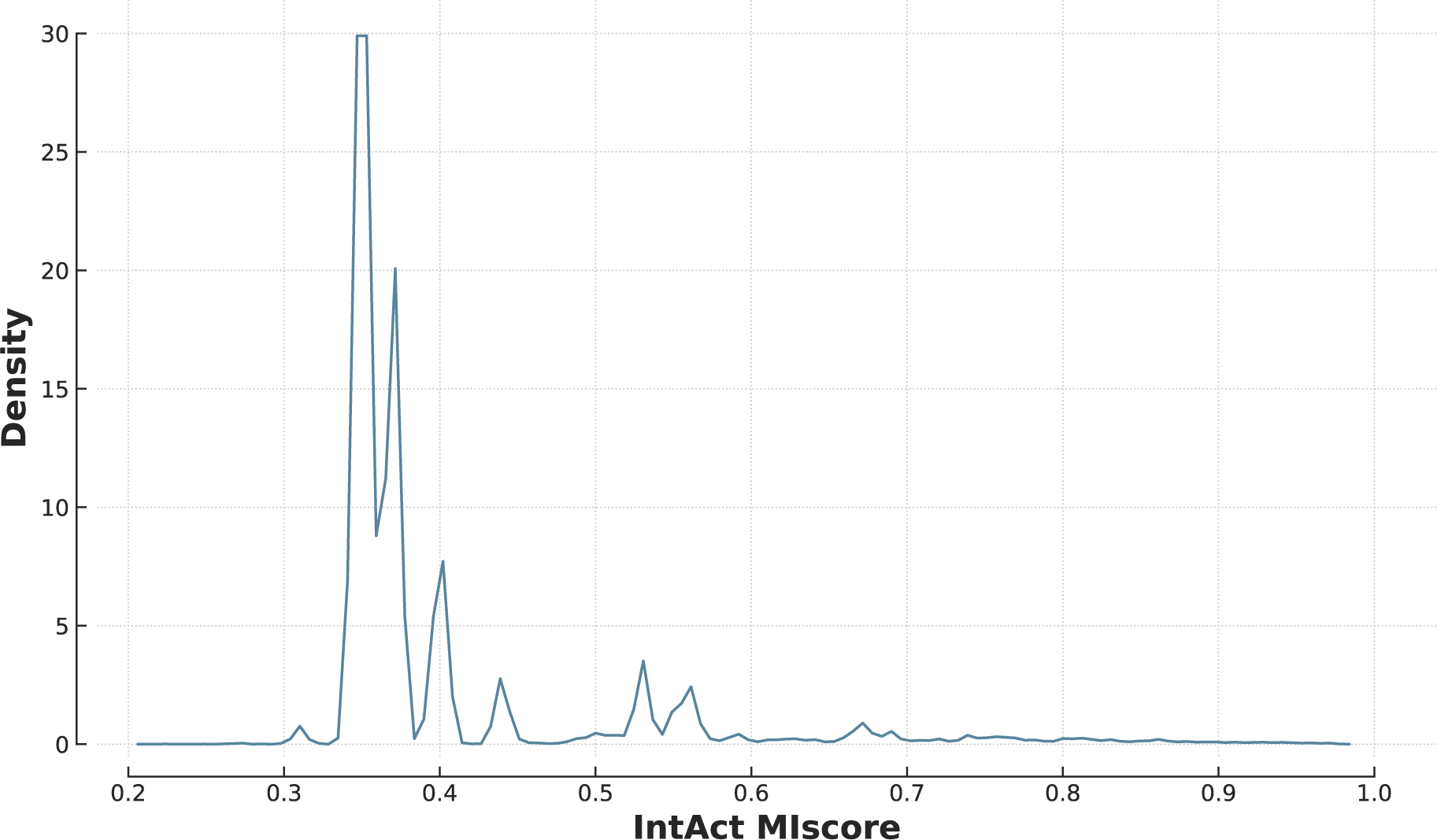
distribution of IntAct’s MIscore in the yeast dataset.

### Supplementary Tables

**Supplementary Table 1:** Sample size in B4PPI.

| Set | Number of examples (% of positive) |  |
| --- | --- | --- |
|  | B4PPI-Human | Yeast dataset |
| Training | 106,662 (50%) | 60,738 (50%) |
| T1 | 24,898 (50%) | 25,398 (50%) |
| T2 | 1,244,900 (1%) | N/A |

**Supplementary Table 2:** Sample size in each category in the dataset used to investigate networks topology.

| Type of interaction | Number of pairs | % of PPIs |
| --- | --- | --- |
| Hub-hub | 27,580 | 50.69 % |
| Hub-lone | 19,205 | 49.70 % |
| Lone-lone | 3,011 | 45.67 % |

**Supplementary Table 3:** Details of the features used for B4PPI-Yeast.

| Feature | Number of different annotations | Missing values (/6,721) | Source |
| --- | --- | --- | --- |
| Biological processes (GO) | 3,114 | 1,510 | UniProt [68] |
| Cellular components (GO) | 820 | 839 | UniProt |
| Molecular functions (GO) | 2,079 | 2,242 | UniProt |
| Domains | 606 | 5,135 | UniProt |
| Motifs | 181 | 6,273 | UniProt |
| Sequence | N/A | 0 | UniProt |

**Supplementary Table 4:**
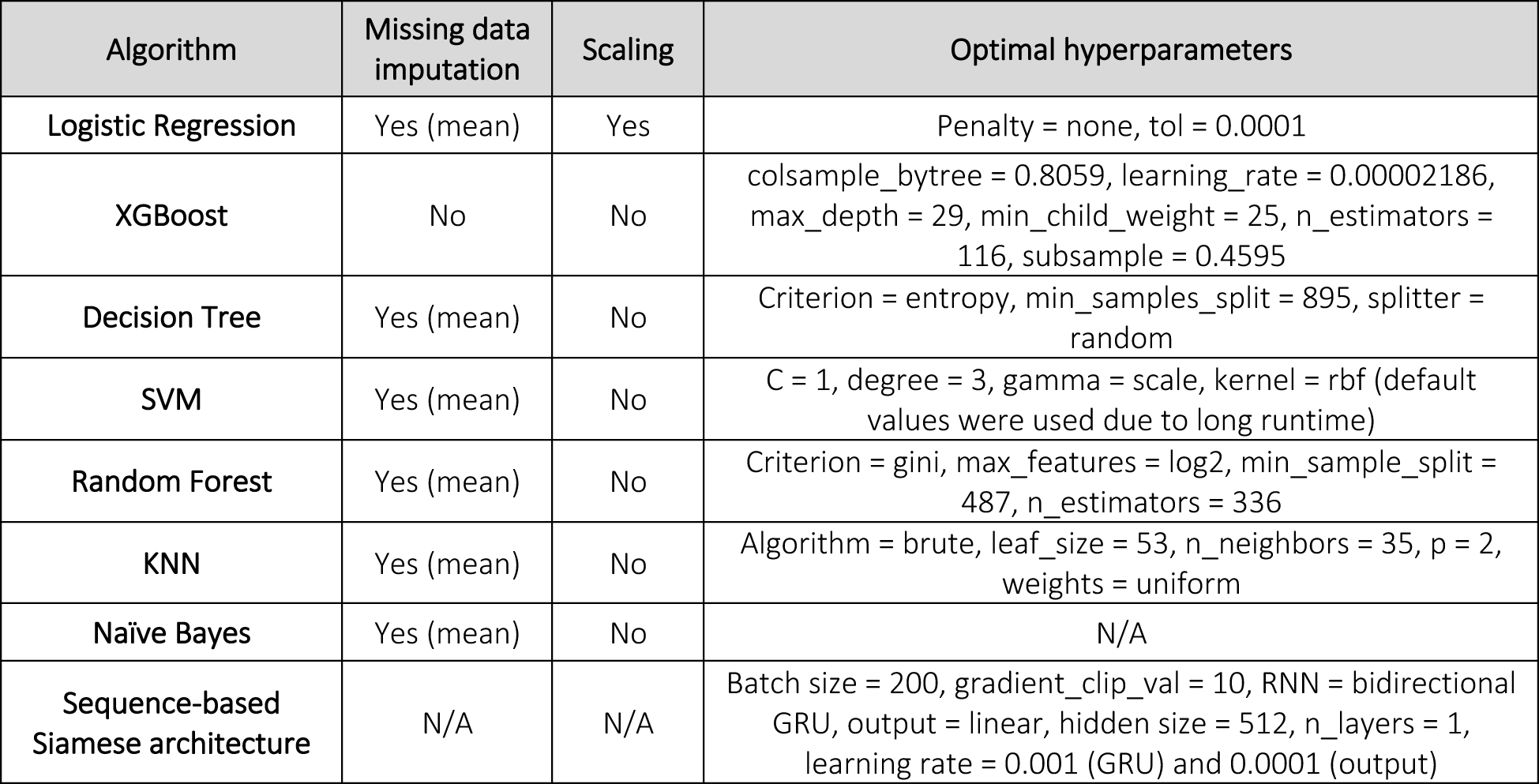
Optimal parameters for the models trained on B4PPI-Human.

| Algorithm | Missing data imputation | Scaling | Optimal hyperparameters |
| --- | --- | --- | --- |
| Logistic Regression | Yes (mean) | Yes | Penalty = none, tol = 0.0001 |
| XGBoost | No | No | colsample_bytree = 0.8059, learning_rate = 0.00002186, max_depth = 29, min_child_weight = 25, n_estimators = 116, subsample = 0.4595 |
| Decision Tree | Yes (mean) | No | Criterion = entropy, min_samples_split = 895, splitter = random |
| SVM | Yes (mean) | No | C = 1, degree = 3, gamma = scale, kernel = rbf (default values were used due to long runtime) |
| Random Forest | Yes (mean) | No | Criterion = gini, max_features = log2, min_sample_split = 487, n_estimators = 336 |
| KNN | Yes (mean) | No | Algorithm = brute, leaf_size = 53, n_neighbors = 35, p = 2, weights = uniform |
| Naïve Bayes | Yes (mean) | No | N/A |
| Sequence-based Siamese architecture | N/A | N/A | Batch size = 200, gradient_clip_val = 10, RNN = bidirectional GRU, output = linear, hidden size = 512, n_layers = 1, learning rate = 0.001 (GRU) and 0.0001 (output) |

**Supplementary Table 5:** Optimal parameters for the models trained on the yeast dataset.

| Algorithm | Missing data imputation | Scaling | Optimal hyperparameters |
| --- | --- | --- | --- |
| Logistic Regression | Yes (mean) | Yes | Penalty = none, tol = 0.0001 |
| XGBoost | No | No | colsample_bytree = 0.7087, learning_rate = 0.00001129, max_depth = 26, min_child_weight = 3, n_estimators = 244, subsample = 0.9318 |
| Naïve Bayes | Yes (mean) | No | N/A |
| Sequence-based Siamese architecture | N/A | N/A | Batch size = 200, gradient_clip_val = 10, RNN = bidirectional GRU, output = linear, hidden size = 512, n_layers = 1, learning rate = 0.001 (GRU) and 0.0001 (output) |

## Notes

https://github.com/Llannelongue/B4PPI

### Competing Interest Statement

The authors have declared no competing interest.

### Summary of Updates

Added analyses on the impact of sequence homology, updated references, updated title

